# The colourless dictyochophyte *Ciliophrys* sp. Baltic secondarily lacks a plastid but hosts a bacterial endosymbiont

**DOI:** 10.64898/2026.03.27.714854

**Authors:** Dovilė Barcytė, Filip Husnik, Marek Eliáš

## Abstract

The loss of an organelle represents an extreme case of reductive evolution. Only a few plastid loss events have been documented and they seem to be particularly rare in free-living ancestrally photosynthetic taxa. We isolated a new phagotrophic protist, *Ciliophrys* sp. Baltic, representing a poorly studied non-photosynthetic lineage in the stramenopile algal class Dictyochophyceae. Genomic and transcriptomic data from this organism demonstrate that it lacks a plastid genome and nuclear genes critical for plastid biogenesis. The plastid organelle is thus most likely secondarily missing from *Ciliophrys* sp. Baltic, although a few enzymes originally targeted to the plastid persist with a changed localization. Unexpectedly, *Ciliophrys* sp. Baltic harbors a bacterial endosymbiont, *Candidatus* Penulousia baltica sp. nov. (Rickettsiales). Analysis of its complete genome sequence uncovered unique aspects of its metabolism not previously reported in other Rickettsiales. The bacterium exploits its host metabolically and presumably manipulates it with an arsenal of putative effector proteins. However, we also found that it might help offset some of the biosynthetic deficiencies of its host stemming from the plastid loss by providing haem and a lysine precursor. Altogether, our study establishes a new single-cell model system for studying plastid loss and endosymbiont acquisition.

## Introduction

Photosynthetic plastids are hallmarks of several major groups of eukaryotes commonly referred to as algae and plants (Adl et al. 2018; Shibbald and Archibald 2020). Notably, nearly all groups of plastid-bearing eukaryotes contain lineages that have transitioned to a non-photosynthetic state by losing components of the photosynthetic machinery (Hadariová et al. 2018; Maciszewski and Karnkowska 2019). The plastid as such has usually been retained in these lineages, a testament to the fact that the key physiological roles of the organelle transcend the emblematic process of photosynthesis. Nevertheless, the secondary absence of a plastid due to its genuine loss has been documented, primarily in parasitic taxa including Syndinea (a group of parasitic dinoflagellates; Gornik et al. 2015; Farhat et al. 2021; Holt et al. 2023) and certain lineages of Apicomplexa (Mathur et al. 2023). However, free-living taxa are not immune to plastid elimination as well. Such elimination may have occurred in Picozoa, a group of tiny free-living heterotrophic flagellates phylogenetically nested within the eukaryotic supergroup Archaeplastida, for which no evidence of a plastid has been found (Schön et al. 2021).

Most recently, plastid loss has been argued to have affected the evolution of the Ochrophyta, a quintessentially algal radiation characterized by the presence of a secondary plastid of red algal origin and including well-known organisms such as kelps and diatoms. Still, secondarily non-photosynthetic members have long been known, being particularly frequent among chrysophytes (Dorrell et al. 2019) and sporadically occurring in other ochrophyte lineages (Sekiguchi et al. 2002; Heesch et al. 2008; Frankovich et al. 2018; Onyshchenko et al. 2019; Kayama et al. 2020; Bringloe et al. 2021; Barcytė et al. 2024). All such cases naturally pose a question of the fate of the plastid organelle, but already early transmission electron microscopy (TEM) studies revealed a candidate residual plastid (often referred to as the leucoplast) in many of these organisms (e.g., Sekiguchi et al. 2002). Genomic evidence further documents the presence of a plastid even in some of the colourless ochrophytes where a candidate plastid organelle could not be discerned by TEM (e.g., Barcytė et al. 2024). Genomic (and transcriptomic) data paint a picture of a high functional variability among ochrophyte leucoplasts, which retain a varying subset of metabolic functions and, in most cases, also their own reduced genome (plastome) and the associated gene expression machinery (Kamikawa et al. 2017; Kayama et al. 2020; Barcytė et al. 2024). Nevertheless, a genome-lacking plastid has been robustly documented from the chrysophyte genus *Paraphysomonas* (Dorrell et al. 2019), marking one of the extremes of the reductive plastid evolution.

This case, however, has been recently surpassed by the first compelling evidence for genuine plastid loss in ochrophytes. Terpis et al. (2025) studied the minute colourless marine protist *Picophagus flagellatus*, representing a separate ochrophyte class (Picophagea). The genomic and transcriptomic data allowed a search for the potential genetic signatures of a plastid, as the previous TEM investigation had not detected the organelle itself (Guillou et al. 1999). Terpis et al. (2025) found that *P. flagellatus* lacks not only a plastome but also all genes for proteins that are assumed to be critical for maintaining a plastid organelle, most importantly including subunits of the complexes mediating protein import of nucleus-encoded proteins into the plastid. In addition, no specific metabolic process that would be compartmentalized in a plastid was identified in *P. flagellatus* based on analyzing its gene repertoire, so the authors concluded that this ochrophyte has undergone complete plastid elimination. The same study and a parallel work by Cho et al. (2024) also showed that the “heliozoan” *Actinophrys sol*, previously demonstrated to lack a plastid (Azuma et al. 2022), is a genuine ochrophyte that evolved from a plastid-bearing ancestor (independently of *P. flagellatus*).

Plastid loss in ochrophytes is not necessarily restricted to *P. flagellatus* and *A. sol*. An additional notable candidate is the genus *Ciliophrys*, a unicellular phagotrophic “helioflagellate” that forms a characteristic Heliozoa-like stage, with the cell decorated by multiple axopodia yet keeping a motile flagellum. Phylogenetic analyses of 18S rRNA gene sequences from multiple *Ciliophrys* isolates revealed pronounced phylogenetic diversity within the *Ciliophrys* morphotype and robustly positioned this lineage into the ochrophyte class Dictyochophyceae, consistent with ultrastructural characteristics. This shallow nesting within the ochrophyte phylogeny makes it practically certain that the *Ciliophrys* lineage has evolved from a plastid-containing ancestor. However, no structure that could be identified as a putative (however reduced) plastid has ever been observed in *Ciliophrys*, and the authenticity of an *rbcL* gene sequence reported earlier from a *Ciliophrys* culture and interpreted as evidence for the presence of a plastid genome (Sekiguchi et al. 2002) was subsequently questioned (Kayama et al. 2020). Hence, the jury is still out on whether *Ciliophrys* possesses a plastid or not.

While plastids (and mitochondria) represent the outcomes of ancient endosymbiotic associations between specific bacteria and eukaryotic hosts, extant eukaryotes continue to provide a niche for more recent endosymbionts. In fact, the recent resurgence in this topic has led to the realization that bacterial endosymbionts are surprisingly diverse and common, occurring across many types of eukaryotic hosts, including protists (Husnik et al. 2021). Especially frequent in protists might be the so-called “professional” endosymbionts, i.e., bacteria specialized for intracellular life, exploitation of their hosts, and horizontal movement between them. Rickettsiales, Holosporales, or Chlamydiae are among the most prominent groups of professional endosymbionts, and their representatives are being reported from members of an ever-growing list of protist lineages (Husnik et al. 2021). In this context, it is notable that encounters of professional endosymbionts in ochrophytes, despite the sheer diversity of this group, have been scarce. We are aware of only two clearly documented cases, both involving members of the family Rickettsiaceae: *Candidatus* Phycorickettsia, which has been reported from multiple members of the ochrophyte class Eustigmatophyceae (Yurchenko et al. 2018), and a representative of the broad “Megaira group” detected in the genome data from the brown alga *Nemacystus decipiens* (Davison et al. 2023).

Here, we revisit the status of the plastid in *Ciliophrys* with genome-scale sequence data from a newly isolated representative of this lineage. Our analyses not only provided compelling evidence for plastid loss in our isolate but unexpectedly also revealed the presence of a bacterial endosymbiont from the “Megaira group”, further broadening the host range of this Rickettsiaceae branch. We report on the complete genome of this bacterium, which provides insights into potential physiological links between the endosymbiont and its host, some of which may compensate for the metabolic capacities lost by *Ciliophrys* sp. Baltic due to the loss of the plastid.

## Results and Discussion

### Ciliophrys sp. Baltic expands the phylogenetic diversity of the non-photosynthetic dictyochophytes

With the aim of obtaining a culture of a *Ciliophrys* representative, we searched for the respective morphotype in natural samples and successfully isolated a candidate from a sand sample collected from a beach on the Baltic Sea in Lithuania. Maximum likelihood (ML) phylogenetic analysis of the 18S rDNA dataset confirmed the morphological identification of our isolate, placing it within a clade of organisms nominally assigned to the genus *Ciliophrys* (Fig. 1; Fig. S1). Specifically, it grouped with an environmental sequence from the Ocean Barcode Atlas (OBA; Vernette et al. 2021), recovered from surface marine water in the opening of the Gulf of Panama, forming a deeply branching lineage within the clade. The substantial genetic diversity within the *Ciliophrys* clade, as documented by the 18S rRNA gene tree, suggests that a more appropriate formal classification would divide its representatives into multiple genera, united within the already established family Ciliophryaceae. For the purpose of this study, we refer to our organism as *Ciliophrys* sp. Baltic and leave detailed taxonomic questions for a future dedicated investigation. The photosynthetic genus *Rhizochromulina* (family Rhizochromulinaceae), which currently includes the single formally described species *Rhizochromulina marina* (Hibberd & Chretiennot-Dinet 1979) but also a diversity of uncharacterized strains nominally assigned to it, was confirmed to be the closest relative of Ciliophryaceae, the two together constituting the order Rhizochromulinales (Fig. 1; Fig. S1). The phylogenetic unity of Rhizochromulinales is consistent with morphological characteristics shared by its representatives including *Ciliophrys* sp. Baltic, such as an amoeboid body form (Fig. 2A), numerous radiating protrusions (Fig. 2B), a flagellum held in a distinctive shape of a loop (Fig. 2C), and fast-swimming spindle-shaped zoospores.

**Fig. 1.**
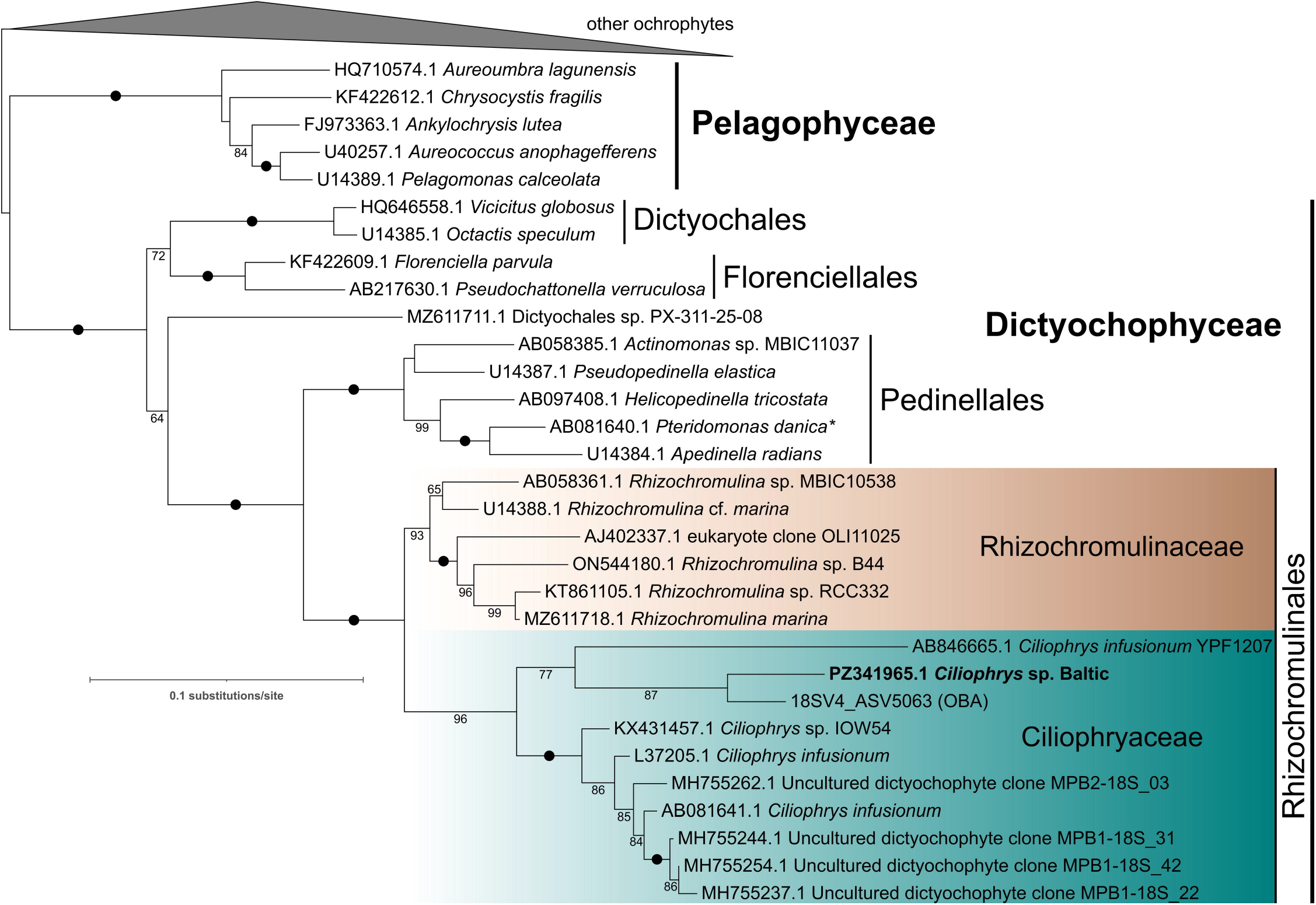
Maximum-likelihood phylogenetic tree of ochrophytes based on 18S rDNA sequences as generated by IQ-TREE. The alignment used for tree inference consisted of 1,867 nucleotide positions, and the substitution model employed was TIM2+F+I+R4. Numbers at branches represent bootstrap values (shown when >50%) calculated from 100 non-parametric bootstrap replicates; black dots mark branches with bootstrap values of 100%. The newly identified sequence of *Ciliophrys* sp. Baltic is shown in bold, the sequence 18SV4_ASV5063 next to it comes from the Ocean Barcode Atlas (OBA; Vernette et al. 2021). The asterisk indicates the colourless relative of *Ciliophrys*, i.e., the genus *Pteridomonas*, which contains a non-photosynthetic plastid. The full version of the tree is presented as Fig. S1.

**Fig. 2.**
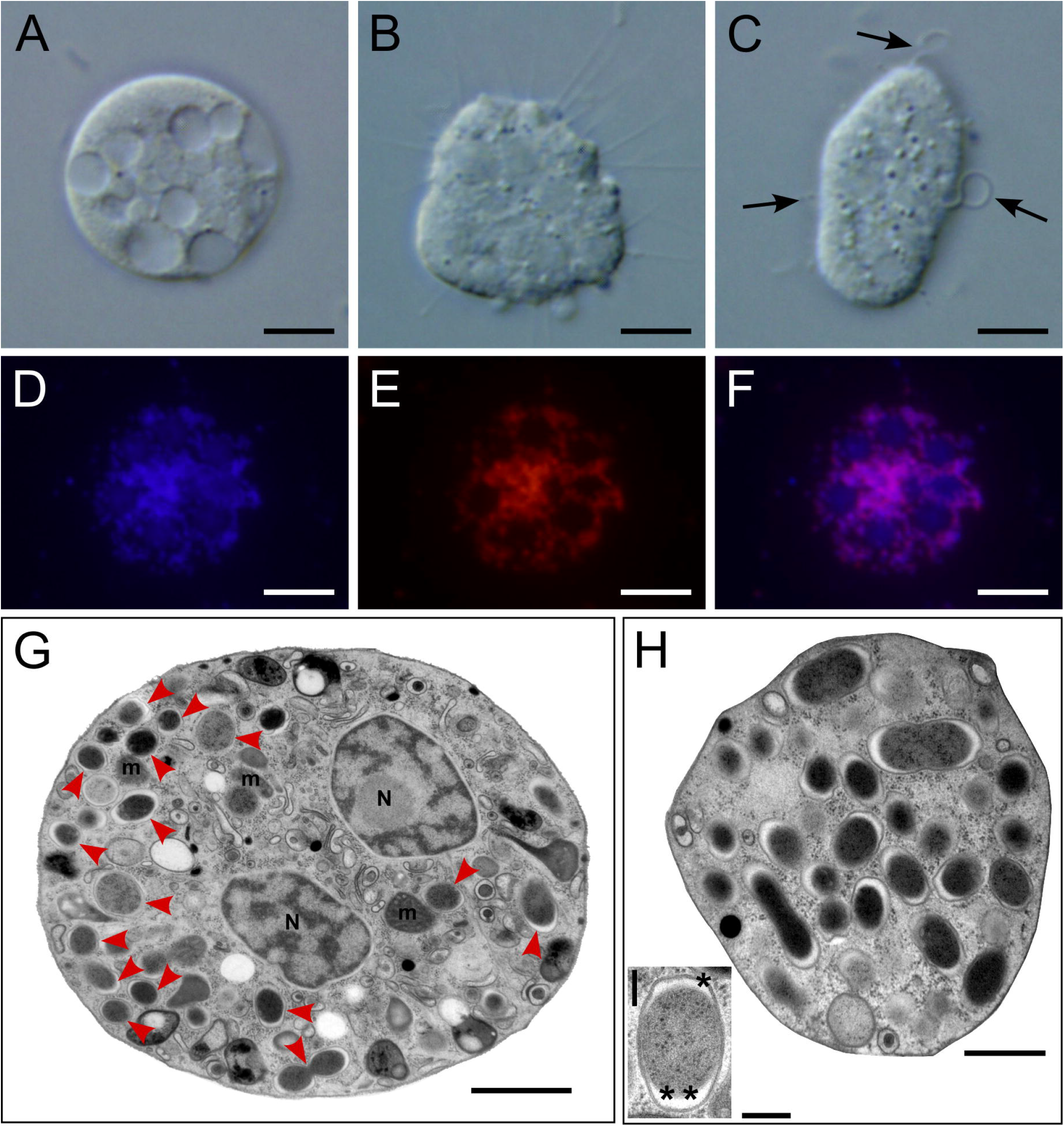
Light, fluorescence, and electron microscopy of *Ciliophrys* sp. Baltic. (A–C) Differential inference contrast micrographs demonstrating a round form of the organism full of vacuoles (A), a multinucleate cell with spiny projections (B), and (C) a ready-to-divide cell with three emergent flagella (arrows). (D–E) Fluorescent micrographs showing (D) DAPI DNA staining, (E) endosymbiont-targeted FISH fluorescence, and (F) overlaid DAPI and FISH images. (G, H) Transmission electron micrographs showing the general view of the cell containing numerous bacterial endosymbionts. (I) A close-up of the bacterium surrounded by two membranes interpreted as the outer membrane (asterisk) and the inner membrane (double asterisk) with the periplasm (inflated at cell poles) in between. Abbreviations: m, mitochondrion; N, nucleus. Arrowheads point to bacterial endosymbionts. Scale bars: 5 µm (A to F), 1 µm (G), 0.5 µm (H), and 0.2 µm (I).

### The plastid genome is missing, and the mitogenome is inflated by a repeat region in Ciliophrys sp. Baltic

To investigate whether *Ciliophrys* sp. Baltic possesses a plastid genome, we subjected it to both Illumina and Nanopore DNA sequencing. A homology search of the two resulting genome assemblies was performed using representative dictyochophyte plastome sequences (NC_043890.1 and NC_056103.1), but no scaffolds or contigs corresponding to a plastid genome were detected. The sequenced plastome sizes of colourless ochrophytes range from 33,539 to 54,809 bp in the dictyochophyte *Pteridomonas* spp. (Kayama et al. 2020), from 53,209 to 62,910 bp in *Spumella*-like chrysophytes (Kim et al. 2020), from 66,116 to 67,763 bp in the diatom *Nitzschia* spp. (Onyshchenko et al. 2019), while *Leucomyxa plasmidifera* PRA-24 (Synchromophyceae) has the plastome size of 44,072 bp (Barcytė et al. 2024). However, despite manually checking scaffolds with a similar size range and inspecting circular-mapping molecules in assembly graphs, no candidate for a plastid genome was found. Notably, we also did not find any candidate for an *rbcL* gene. Sekiguchi et al. (2002) reported an *rbcL* sequence from a distantly related *C. infusionum* strain (sequence AB081641.1 in Fig. 1; Fig. S1), which they interpreted as evidence for the presence of a cryptic vestigial plastid containing a genome and a functional RuBisCO enzyme involved in photosynthesis-unrelated processes. However, a more recent study has shown that this putative *C. infusionum rbcL* sequence is highly similar to that of a photosynthetic Rhizochromulinaceae species (Kayama et al. 2020). Combined with our results, the sequence reported by Sekiguchi et al. is best interpreted as a contaminant.

The absence of plastome sequences in our genomic data from the *Ciliophrys* sp. Baltic is unlikely to be due to insufficient sequencing depth, as demonstrated by the fact that we readily obtained the mitochondrial genome (mitogenome) of the organism, assembled as a circular-mapping molecule of 42,318 bp. A total of 60 standard mitochondrial genes, including 33 genes encoding conserved mitochondrial proteins (counting the split *cox2* gene as one gene; see below), 24 tRNA genes, and three rRNA genes, were annotated in it (Fig. 3). In terms of the gene content, the *Ciliophrys* sp. Baltic mitogenome did not reveal any unexpected features, with all the genes being shared with at least some previously investigated ochrophytes (Ševčíková et al. 2016; Sibbald et al. 2021; Barcytė et al. 2024). One of the non-conserved ORFs, *orf211*, occurs in the genomic context (directly downstream of the *rpl2-rps19-rps3-rpl16* gene block) where the *rps1* gene was recognized in mitogenomes of the related class Pelagophyceae (Sibbald et al. 2021). We tested the possibility that *orf211* represents an *rps1* ortholog by a HHpred search, which did not return any significant hit when the *orf211-*encoded protein sequence was used as a query. Likewise, no significant hits were found when the protein’s tertiary structure was predicted with two different tools and compared against databases of experimentally determined or predicted protein structures. Hence, if *orf211* is orthologous to *rps1*, it has diverged beyond recognition. It is notable that *rps1* was until recently known to occur only in pelagophytes among all stramenopile groups (Sibbald et al. 2021), but recently an extremely divergent version was identified in the mitogenomes of the class Eustigmatophyceae (Richtář et al. 2026), suggesting that the gene may generally be rapidly evolving in ochrophyte mitochondria and, therefore, difficult to recognize. The remaining three non-conserved ORFs are contained in two introns interrupting the *cox1* gene, with two of them (*orf511* and *orf552*) encoding putative reverse transcriptase/maturase proteins. The *cox2* gene is split into what are apparently two separate exons, arranged in the genome as a tandem in an order opposite to how they are expected to appear in a complete transcript, and thus most likely reconstituting a mature mRNA via *trans*-splicing. The specific set of tRNA genes implies the import of at least two tRNAs from the cytosol to meet the needs of mitochondrial translation, namely tRNAs decoding threonine codons (as in stramenopiles in general; Burger and Nedelcu 2012) and the CGN box of arginine codons.

**Fig. 3.**
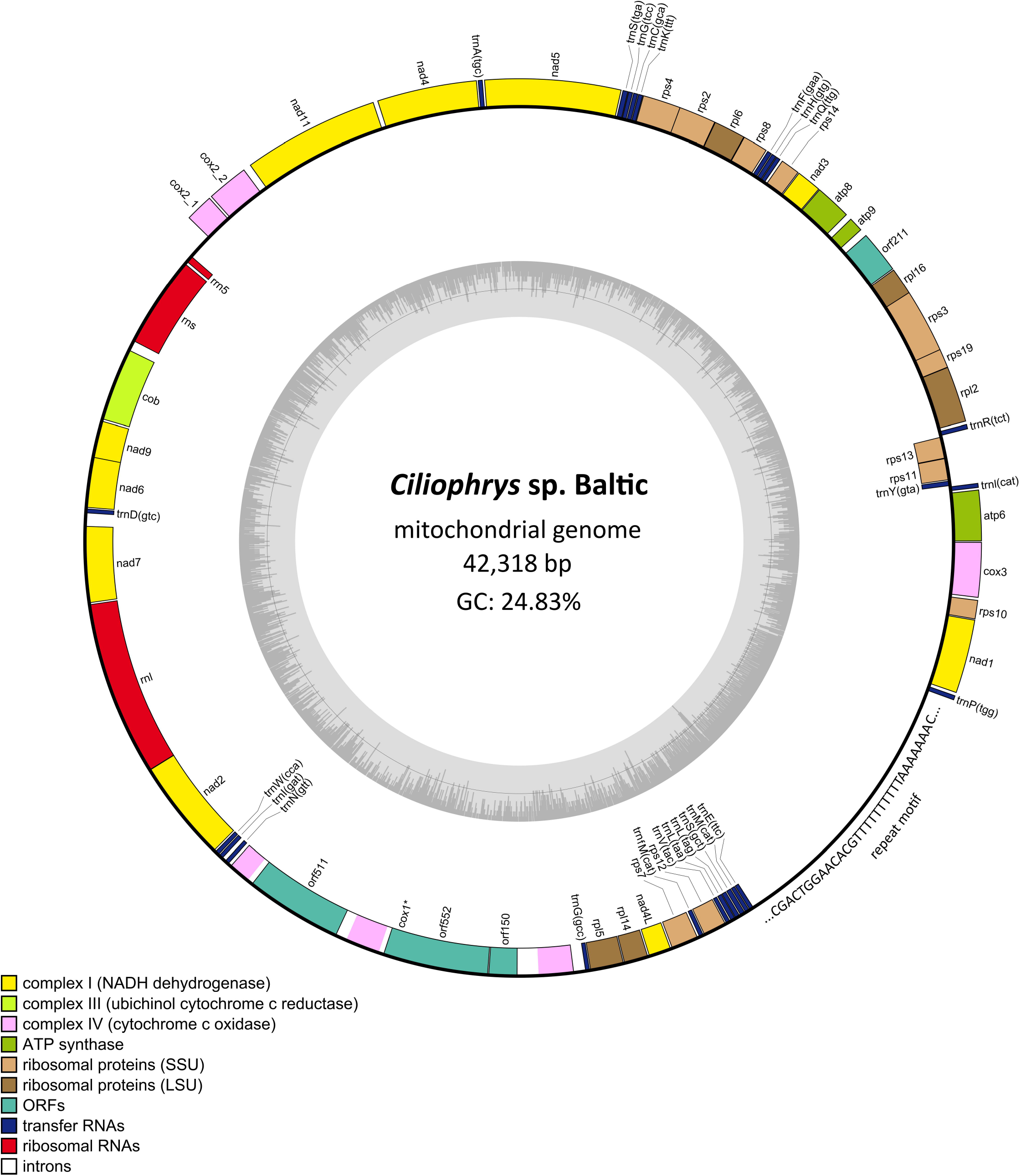
Gene map of the mitochondrial genome of *Ciliophrys* sp. Baltic. Genes are represented as blocks, with different colours indicating the functional categories of the genes. The inner circle plot displays the GC content, with the thin grey line marking 50%. The *cox1* gene (asterisk) is interrupted by two introns: one with a single ORF (*orf511*) encoding a putative reverse transcriptase/maturase, and another with two ORFs, one of which (*orf552*) also encodes a putative reverse transcriptase/maturase. A repeat motif is shown next to the tandem repeat region.

Notably, the mitogenome of *Ciliophrys* sp. Baltic contains two blocks of tandem repeats confined to a single large region lacking any annotated genes, spanning 4,382 bp. Both blocks are composed of the same repeat motif, yet on the opposite strands, and are separated by a unique region of ∼200 bp. The repeated sequence unit CGACTGGAACACGTTTTTTTTTTAAAAAAAC, when up to two mismatches are allowed for, is present 95 times in the first block (or 100 times if three mismatches are permitted) and 30 times (31 times with the more relaxed definition) in the second block. Analogous large tandem repeat regions have also been found in mitogenomes of other ochrophytes, including the diatom *Phaeodactylum tricornutum* (Oudot-Le Secq & Green 2011) and the pelagophyte *Aureoumbra lagunensis* (Sibbald et al. 2021), as well as in haptophytes such as *Chrysochromulina tobinii* (Hovde et al. 2014) and *Isochrysis galbana* (Fang et al. 2022). This region may form an unusual structure in the *Ciliophrys* sp. Baltic mitochondrial DNA, but its function, if any, is unclear.

### Ciliophrys sp. Baltic lacks a plastid yet exhibits its genetic footprints

The results presented above allowed us to conclude that *Ciliophrys* sp. Baltic represents a rare case of an ochrophyte that has secondarily lost the plastid genome. This is, however, not a definite indicator of the disappearance of the organelle itself. A case in point is the chrysophyte genus *Paraphysomonas*, which retains the cryptic organelle without a genome (Dorrell et al. 2019). Genome-lacking plastids are also known to occur in various protist and plant lineages outside of ochrophytes (Maciszewski and Karnkowska 2019; Gawryluk et al. 2019). We therefore set out to evaluate the presence of plastid-targeted proteins in our organism using a transcriptome assembly. According to a BUSCO analysis, its completeness was estimated to be 80% (using the stramenopiles_odb10 dataset as a reference). As a positive control, we searched for a set of nucleus-encoded proteins serving as hallmarks of the mitochondrion and previously used for the same purpose (Terpis et al. 2025). All of them were identified (Table S1), except for homologs of Min proteins, which are not universally present in ochrophytes (Leger et al. 2015); notably, we did not detect them even in the *Ciliophrys* sp. Baltic genome assembly. We thus conclude that the transcriptome assembly captures a substantial and representative portion of the expected gene repertoire and is suitable for downstream analyses. In parallel, we also examined our draft genome assembly. Our strategy to evaluate the presence of genomic signatures of a plastid organelle essentially followed that employed by Barcytė et al. (2024) and Terpis et al. (2025).

First of all, if physically present, the plastid would contain protein complexes that mediate the import of nucleus-encoded proteins, which must traverse the multiple membranes constituting the plastid envelope, expected to be four as in other ochrophytes. Therefore, we searched our transcriptome and draft genome assemblies for homologs of SELMA (symbiont-specific ERAD-like machinery) as well as TOC and TIC components (translocons of the outer and inner chloroplast membranes, respectively) (Maier et al. 2015). We detected two proteins containing Der1/Der2-like family domains (Table S1); however, they obviously represent components of the host ERAD machinery (Fig. S2). Meanwhile, no candidates for TOC and TIC components were present, and the stromal processing peptidase (an enzyme responsible for removing the transit peptide of proteins upon their import into the stroma) was likewise not identified. We also searched for homologs of the core components of the plastid division system in ochrophytes, i.e., plastidial FtsZ and the dynamin-related protein DRP5B (also known as ARC5; Miyagishima et al. 2014), but detected only the mitochondrial version of the former (Table S1) and dynamin-related proteins non-orthologous to DRP5B. For comparison, homologs of the proteins critical for plastid maintenance were identified in transcriptome assemblies from other (i.e., plastid-bearing) dictyochophytes including the non-photosynthetic *Pteridomonas danica*, and a subset of them also occurs in the chrysophyte *Paraphysomonas imperforata* that possesses a genome-lacking leucoplast (Table S2).

As a further test, we searched for a periplastidial compartment (PPC)-targeted Hsp70 isoform involved in chaperoning plastid preproteins to the Tic/Toc import apparatus (May and Soll 2000). Seven nucleus-encoded Hsp70 candidates were detected in *Ciliophrys* sp. Baltic after excluding obvious bacterial contaminants (Table S1). Four of them lacked an N-terminal extension, two contained a signal peptide, and the last one possessed a predicted mitochondrial transfer peptide. Phylogenetic analysis assigned these seven proteins into four subgroups (Fig. S3). Three of the subgroups represented well-established cytosolic, endoplasmic reticulum-localized, and mitochondrial Hsp70 versions, and the assignment of particular *Ciliophrys* sp. Baltic proteins into them was fully consistent with the results of subcellular targeting predictions. One *Ciliophrys* sp. Baltic protein, together with orthologs from other stramenopiles, fell into a potentially novel subgroup of unknown function, but presumably consisting of cytosolic proteins owing to the absence of any N-terminal presequence. Crucially, the clade containing proteins with experimentally confirmed PPC localization (Gould et al. 2006; Gruber et al. 2025) did not include any *Ciliophrys* sp. Baltic homolog. Finally, we checked two enzymes, aspartate aminotransferase and phosphoenolpyruvate carboxylase, for which PPC-targeted versions exists in model diatoms (Tanaka et al. 2014; Zhang et al. 2025). However, the only homologs detected in *Ciliophrys* sp. Baltic were more similar to the mitochondrial versions of the enzymes in the respective diatom species and were likewise predicted to bear N-terminal mitochondrial transfer peptides (Table S1).

These findings indicate the absence of a plastid in *Ciliophrys* sp. Baltic, a notion further supported by our *in silico* reconstruction of its metabolic repertoire. The first pathway we examined was haem biosynthesis, in ochrophytes taking place exclusively in plastids (Cihlář et al. 2019). However, we found no evidence of haem biosynthesis enzymes being present in *Ciliophrys* sp. Baltic, suggesting that the organism relies on external haem sources. The presence of a mitogenome encoding components of the respiratory complexes (see above) makes it clear that *Ciliophrys* sp. Baltic requires haem. In addition, homologs of the two enzymes catalyzing the conversion of protohaem to haem A, an essential component of the respiratory complex IV, i.e. COX10 and COX15, are found in the *Ciliophrys* sp. Baltic transcriptome assembly (Table S1). Thus, the question arises as to how (proto)haem supply to *Ciliophrys* sp. Baltic is secured. The diet is an obvious possibility, but an alternative source is suggested by an unexpected finding presented below. Like *Ciliophrys* sp. Baltic, the previously identified aplastidic ochrophytes (*P. flagellatus* and *A. sol*) also lack all haem biosynthesis enzymes despite requiring this molecule for their metabolism (Azuma et al. 2020; Terpis et al. 2025). The isoprenoid precursors can be produced by two alternative routes, the mevalonate (MVA) or the non-mevalonate (MEP/DOX) pathway. In eukaryotes, the former typically takes place in the cytoplasm, whereas the latter is exclusively localized in plastid organelles, including the non-photosynthetic ones (Guggisberg et al. 2014). We found no evidence of the MEP/DOX pathway in *Ciliophrys* sp. Baltic, while a full set of MVA pathway enzymes is encoded by the organism (Table S1).

The type II fatty acid synthase (FASII) system is the principal pathway for *de novo* fatty acid biosynthesis in plastid-bearing eukaryotes and is localized to the plastid (Ryall et al. 2003; Kohli et al. 2016). Searches for homologs of the six enzymes of this pathway in the transcriptome and genome data from *Ciliophrys* sp. Baltic yielded outcomes of three different types (Table S1). Two of the enzymes retrieved no homologs. Two other enzymes do have homologs in *Ciliophrys* sp. Baltic, but those represent non-orthologous versions, specifically isoforms that in plastid-bearing dictyochophytes and other ochrophytes co-occur with the plastid-targeted versions yet lack N-terminal presequences. Only the two remaining enzymes (FabD and FabI) are represented in *Ciliophrys* sp. Baltic by putative orthologs of the plastid-targeted versions. However, none of these proteins grouped with their expected closest relatives, i.e., FabD and FabI proteins from dictyochophytes, in the inferred phylogenies (Figs. S4, S5), possibly due to an accelerated sequence evolution upon plastid loss. In the case of FabD, the tree inference resulted in a clade (unsupported by bootstrap analysis) that combines sequences from all three unrelated plastid-lacking ochrophytes (*Ciliophrys* sp. Baltic, *A. sol*, and *P. flagellates*; Figs. S4). Crucially, the N-terminal regions of both FabD and FabI in *Ciliophrys* sp. Baltic have been remodelled and instead of bearing resemblance to the original plastid-targeted presequences, they are in both cases predicted to serve as mitochondrial transfer peptides (Table S1; Fig. S6). Hence, four of the ancestral genes of the plastidial FASII pathway were lost in the *Ciliophrys* lineage, and only two have persisted as genetic footprints of the vanished plastid organelle, encoding proteins retargeted to a different compartment. It is possible that *Ciliophrys* sp. Baltic employs these enzymes in the production of lipoic acid in the mitochondrion. It is interesting to note that *Ciliophrys* sp. Baltic does not contain type I FAS, unlike *P. flagellatus* and certain related ochrophytes, which seems to have acquired it via horizontal gene transfer (HGT) (Terpis et al. 2025).

Enzymes of the plastid-associated shikimate pathway and its branches leading towards aromatic amino acids (phenylalanine, tyrosine, and tryptophan) and folate, are also missing in *Ciliophrys* sp. Baltic. While this organism does encode enzymes for assimilative reduction of sulfate to sulfite (PAPSS and CysH; Table S1), it lacks sulfite reductase, a common plastid-localized enzyme generating H_2_S for cyteine synthesis, as well as any candidate for plastidial version of cysteine synthase (CysK) that assimilate the H_2_S to yield cysteine in the organelle. The three CysK homologs identified in *Ciliophrys* sp. Baltic are predicted to be targeted to the mitochondrion or to possess and N-terminal transmembrane segment (Table S1), and each is more similar to a different non-plastidial CysK isoform in other dictyochophytes than to their plastid-targeted version (e.g., HBNI01012318.1 from *Florenciella parvula*). Lysine biosynthesis is another metabolic process confined to the plastid in ochrophytes (Dorrell et al. 2017), but *Ciliophrys* sp. Baltic recapitulates the situation previously encountered in *P. flagellatus* (Terpis et al. 2025) in that *de novo* lysine biosynthesis is missing and only homologs of the final two enzymes of the pathway (diaminopimelate epimerase and diaminopimelate decarboxylase) are present (Table S1). Crucially, both lack an N-terminal presequence, indicating they are most likely localized to the cytosol (Fig. S6). The biosynthesis pathways of riboflavin and vitamin E (tocopherols and tocotrienols) are also found in plastids, including those of at least some non-photosynthetic ochrophytes (Dorrell et al. 2019; Barcytė et al. 2024), but our data indicate their absence in *Ciliophrys* sp. Baltic, further strengthening the case for the complete elimination of the plastid organelle in this specific organism.

Finally, we investigated whether *Ciliophrys* sp. Baltic encodes the pathways for the synthesis of the characteristic structural lipids of plastid membranes (Yoshihara & Kobayashi 2022). Homologs of the enzymes responsible for the production of galactolipids, i.e. monogalactosyldiacylglycerol (MGDG) and digalactosyldiacylglycerol (DGDG), are lacking in this organism, and while the two enzymes for the synthesis of phosphatidylglycerol are present, closer scrutiny revealed they correspond to the commonly occurring mitochondrial versions (Table S1). Hence, as a striking exception to the pattern described above, came the identification of a complete set of enzymes required for the synthesis of the sulfolipid sulfoquinovosyldiacylglycerol (SQDG), a structural lipid enriched in thylakoid membranes and not known from any aplastidic eukaryote. Specifically, *Ciliophrys* sp. Baltic encodes homologs of the enzymes mediating assimilative reduction of sulfate to sulfite, as well as of UDP-sulfoquinovose synthase (SQD1) that catalyses a reaction between UDP-glucose and sulfite to yield UDP-sulfoquinovose. The latter compound is a substrate of the final enzyme of the pathway sulfoquinovosyltransferase (SQD2), likewise conserved in *Ciliophrys* sp. Baltic, which completes the synthesis of SQDG by transferring the sulfoquinovosyl moiety onto diacylglycerol. Both UDP-glucose and diacylglycerol are common metabolic intermediates whose provision in *Ciliophrys* sp. Baltic is unsurprisingly secured by standard enzymes (Table S1). Hence, *Ciliophrys* sp. Baltic apparently synthesizes SQDG, but the physiological role of this compound is unclear.

To understand the subcellular localization of the SQDG biosynthesis in *Ciliophrys* sp. Baltic we compared the N-terminal sequences of the respective enzymes with their homologs from other, i.e. plastid-containing, dictyochophytes (Fig. S7). Both SQD1 and CysH, i.e. phosphoadenosine phosphosulfate reductase providing sulfite for SQD1, are expected to be localized in the plastid stroma, and hence to exhibit typical plastid-targeting presequences including a signal peptide directly followed by a transit peptide-like region starting with a conserved phenylalanine residue critical for targeting to the protein into the plastid stroma (Gruber et al. 2025). The proteins from plastid-bearing dictyochophytes indeed all fit this expectation. In contrast, while the *Ciliophrys* sp. Baltic orthologs of both proteins contain a robustly predicted signal peptide, in CysH it is directly followed by a sequence that corresponds to the mature protein (the enzyme itself), whereas in SQD1 the region where a transit peptide is expected is shorter and exhibits a markedly different amino acid composition compared to the transit peptide-like region in the other dictyochophyte homologs, and lacks the functionally critical phenylalanine residue or its less common alternative tryptophan (Fig. S7).

SQD2 in plant plastids is associated with the inner envelope membrane, presumably facing the stromal side (Tietje & Heinz 1998). Whether this is true also for SQD2 in dictyochophytes is not certain, because the N-terminal region of SQD2 proteins from plastid-bearing dictyochophytes is more variable in sequence and length than the N-terminus of SQD1 and CysH, the prediction of the expected signal peptide is less consistent, and the amino acid residue following the predicted cleavage site is generally not phenylalanine. These observations indicate that the protein is not targeted to the stroma or reaches the compartment in a non-canonical way. The features of SQD2 from *Ciliophrys* sp. Baltic are similar (Fig. S7), so the only conclusion we can presently draw on its subcellular localization is that like SQD1 and CysH in this organism, it most likely enters the ER lumen in a signal peptide-mediated manner. We may only speculate what happens next with these proteins, but an interesting possibility to ponder is that they are trafficked into a specialized compartment of the endomembrane system that has persisted as a remnant after the plastid itself (i.e. the structure delimited by the membranes containing the SELMA, TOC and TIC complexes) has been lost. Furthermore, this hypothetical compartment is distinct from the endoplasmic reticulum (ER), as none of the three proteins contain the characteristic C-terminal ER retention signal ([KH]DEL).

### Ciliophrys sp. Baltic harbours a Rickettsiaceae endosymbiont, Candidatus Penulousia baltica sp. nov

*Ciliophrys* sp. Baltic feeds on bacteria, and our analysis of the metagenome data from the *Ciliophrys* sp. Baltic culture revealed the presence of a taxonomically diverse bacterial community (Fig. S8). Apart from various bacteria assumed to be free-living based on their taxonomic identification, the community also included a member of the alphaproteobacterial family Rickettsiaceae known to consist of obligate predominantly intracellular symbionts of diverse eukaryotic hosts, including both heterotrophic and phototrophic protists (Husnik et al. 2021). Phylogenetic analysis employing the 16S rRNA gene sequences revealed that the Rickettsiaceae bacterium belongs to the broader “Megaira group”, specifically its internal Clade B as defined by Lanzoni et al. 2019 (Fig. 4; Fig. S9). Using a fluorescently labelled oligonucleotide probe (CilioEndo16S) we found that the bacterium exclusively resided within *Ciliophrys* sp. Baltic cells. The intracellular bacteria densely populated the entire host cell, as evidenced by the widespread fluorescence signal (Fig. 2D–F). Numerous bacteria scattered irregularly throughout the host cytoplast were also visualized by TEM (Fig. 2E, G). They were abundantly present in every *Ciliophrys* sp. Baltic cell observed with TEM and resided directly within the host cytoplasm, i.e., they were not enclosed by any host-derived membrane-bound or peribacterial vacuoles (Fig. 2I). The bacteria were oval-shaped, measuring 0.3–0.6 µm in length and 0.2–0.35 µm in width. Elongated forms, possibly dividing cells reaching up to 0.8 µm in length, were also observed.

**Fig. 4.**
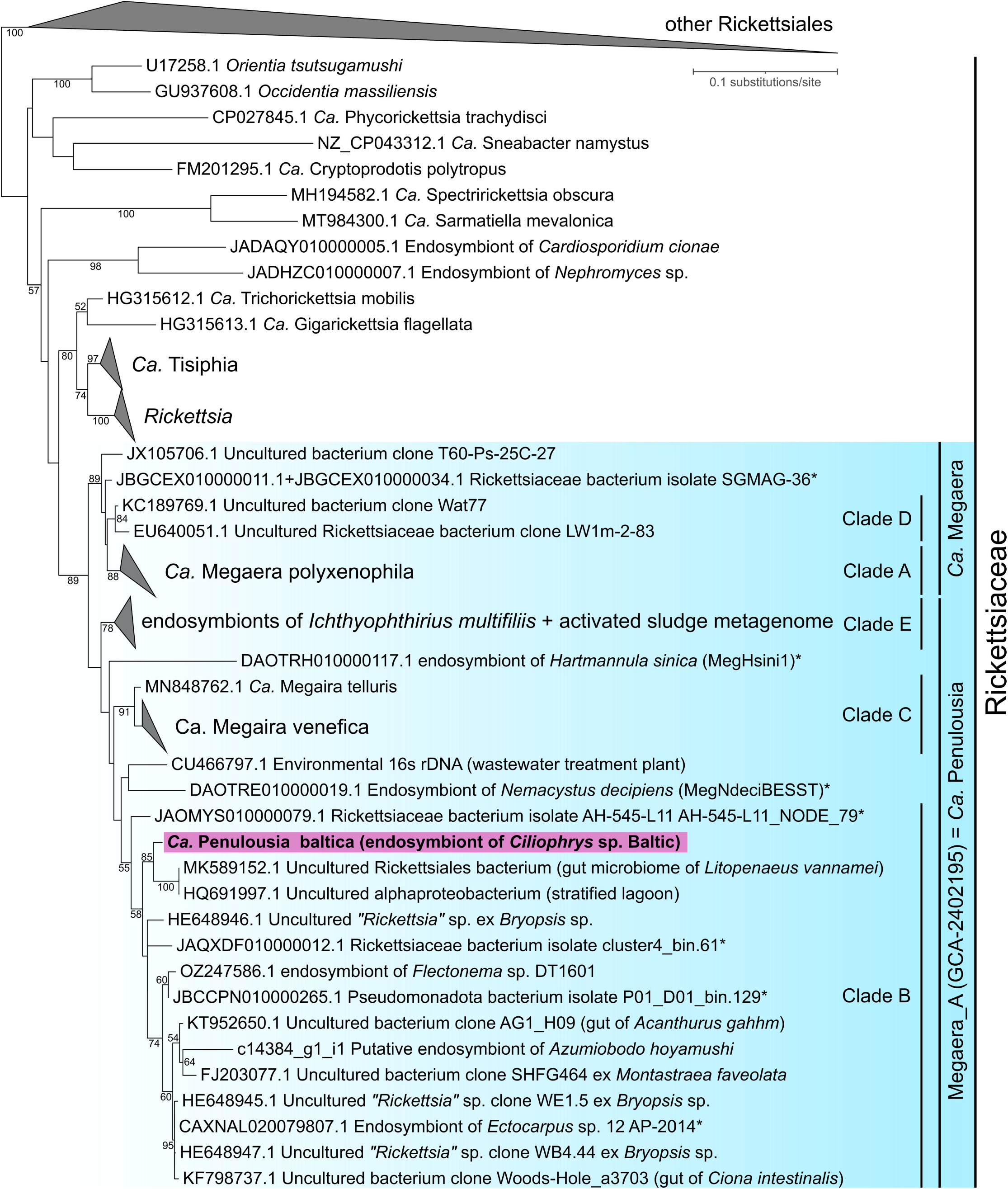
Phylogenetic position of the novel Rickettsiales bacterium from *Ciliophrys* sp. Baltic based on 16S rDNA sequences. The tree was inferred with IQ-TREE from an alignment comprising 1,503 positions with the GTR+F+R6 substitution model. Numbers at branches (shown when >50%) represent bootstrap values (calculated from 100 non-parametric bootstrap replicates). The analysis focuses on the “Megaira group” (highlighted in turquoise). The designation of Clades B to E within it follows the convention introduced by Lanzoni et al. (2019). Some members of the “Megaira group” do not fit into the previously established clades, as already noted by Davison et al. (2023). Note that the “Megaira group” is divided (though with little statistical support) into two subclades that equivalent to two separate genera (*Ca*. Megaera and Megaera_A (GCA-2402195)/*Ca*. Penulousia) strongly supported by a phylogenomic analysis (see Fig. 5). Asterisks indicate MAGs included in the phylogenomic analyses. The full version of the tree is presented as Fig. S9.

To taxonomically name the *Ciliophrys* sp. Baltic endosymbiont, we must address the unsettled taxonomy of the “Megaira group”. An alternative (corrected) spelling of the genus name, “*Megaera*”, was proposed (Oren et al. 2020) and adopted by NCBI, the Genome Taxonomy Database (GTDB; Parks et al. 2022), and some of the recent papers (Giovannini et al. 2026), although not others (Davison et al. 2023; George et al. 2023). More importantly, high genomic diversity within the “Megaira group” was noticed before, supporting the split of the clade into multiple genera (Davison et al. 2023). The objective, relative evolutionary divergence-based rank-normalized classification employed by GTDB divides the representatives of the clade into two genera, *Megaera* (including *Megaera polyxenophila*) and a genus with the placeholder name Megaera_A (GTDB release 11-RS232; GCA-2402195 in the previous releases), which also includes the endosymbionts belonging to the Clade B. This split of the “Megaira group” into two monophyletic subgroups equivalent to *Megaera* and Megaera_A is also supported by our phylogenetic analyses (Fig. 5). Crucially, the formal name *Candidatus* Penulousia has been proposed as an equivalent of Megaera_A (or GCA-2402195; Pallen et al. 2022), which we accept here. However, given the novelty of our discovered bacterium, we refer to it as *Ca. Penulousia baltica* sp. nov. (hereafter *P. baltica*), with the species epithet referring to the geographical origin of the type biological material.

**Fig. 5.**
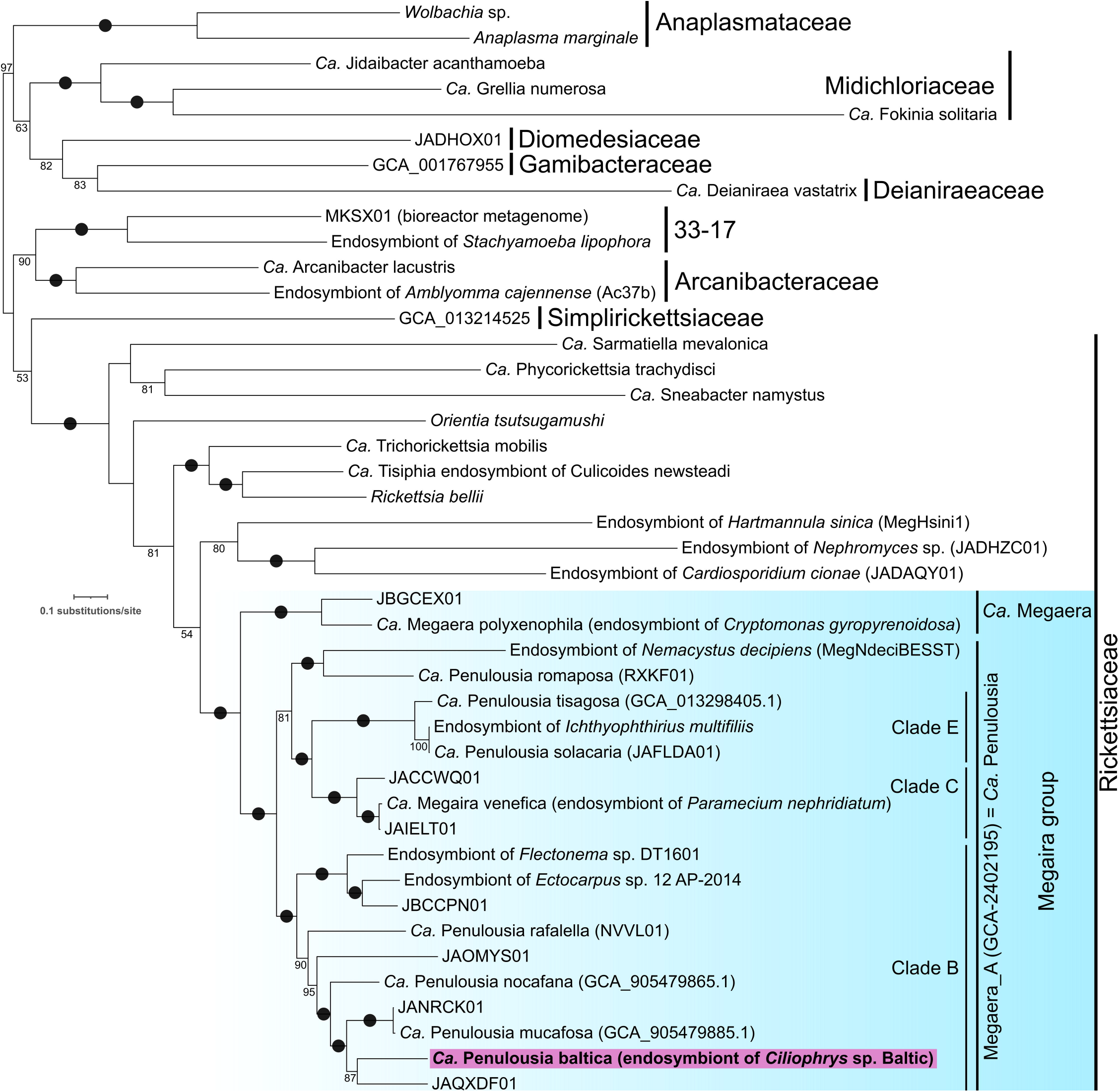
Phylogeny of crown Rickettsiales based on 45 orthologous protein sequences. The tree was inferred with IQ-TREE from a dataset comprising 15,893 aligned positions with the LG+F+I+G4 substitution model. Numbers at branches (shown when >50%) represent bootstrap values (calculated from 100 non-parametric bootstrap replicates); black dots mark branches with bootstrap values of 100%. The new sequence is highlighted. Labels without organism names represent previously assembled environmental MAGs with unidentified host organisms. The division of the “Megaira group” into the separate taxa (genera) *Candidatus* Megaera and Megaera_A (GCA-2402195), i.e., *Candidatus* Penulousia, follows the current release of the Genome Taxonomy Database. Note that the presence of *Ca*. Penulousia endosymbionts in the brown alga *Ectocarpus* sp. 12 AP-2014 and the diplonemid *Flectonema* sp. DT1601 has not been previously documented and is here inferred only from the sequence data available in GenBank.

Based on the 16S rRNA phylogeny, the closest relative of *P. baltica* is represented by the two identical sequences from unknown hosts (obtained from the gut microbiome of a prawn and an environmental sample from a lagoon). Other Clade B members include bacteria found in a green alga (*Bryopsis* sp.), a brown alga (*Ectocarpus* sp.), and a diplonemid (*Flectonema* sp.). Furthermore, we identified a Clade B 16S rRNA gene sequence in a transcriptome assembly from the kinetoplastid *Azumiobodo hoyamushi* (Fig. 4; Fig. S9). Hence, the genus *Ca*. Penulousia (even in its possible narrower concept, equivalent to Clade B) exhibits the potential to associate with phylogenetically diverse hosts. How commonly species of *Ca*. Penulousia may occur in representatives of the class Dictyochophyceae (beyond *Ciliophrys* sp. Baltic) is to be investigated properly, but the only previously reported candidate case seems to be the observation of bacterial endosymbionts of an unknown taxonomic identity in the likewise non-photosynthetic species *Pt. danica* (Patterson & Fenchel 1985). However, we did not identify any Rickettsiales-derived 16S rRNA sequences in any of the multiple dictyochophyte transcriptome assemblies presently available in the NCBI TSA, including an assembly from *Pt. danica* (GenBank HBOW00000000.1; it is not clear whether the sequenced strain corresponds to the isolate reported to harbour endosymbionts). As the *P. baltica* 16S rRNA sequence is abundantly represented in our transcriptome assembly from *Ciliophrys* sp. Baltic, we infer that the other sequenced dictyochophytes are devoid of Rickettsiales endosymbionts.

### Does the endosymbiont provide essential metabolites to its Ciliophrys sp. Baltic host?

With the help of long sequencing reads, we obtained a complete sequence of genome of *P. baltica* and analysed it to gain additional insights into the biology of the bacterium. The genome was assembled as a circular-mapping chromosome of 1,513,035 bp with the GC content of 35.38% (Fig. S10). Using the Rickettsiales dataset as a reference, the assembled genome was determined to be 98.9% complete in BUSCO, and we therefore consider it complete. No plasmids that would possibly belong to *P. baltica* were identified in the genomic data determined from the *Ciliophrys* sp. Baltic culture. Our genome annotation pipeline predicted in the *P. baltica* genome 1,329 protein-coding genes, 34 tRNA genes, a single tmRNA gene, and 3 rRNA genes. The latter are unlinked (the 16S rRNA gene is present elsewhere in the genome than the 23S-5S rRNA gene block) instead of forming a conventional single operon (Fig. S6). This type of arrangement has been documented in other Rickettsiales and is proposed to result from genome degradation (Brewer et al. 2019), a common feature of endosymbiotic bacteria manifested also by gene loss and accumulation of pseudogenes. While a number of such candidates were identified in the *P. baltica* genome using Pseudofinder, our manual curation confirmed the pseudogene status for only a small number of them, mostly representing degraded versions of genes corresponding to mobile genetic elements and duplicated fragments of genes that are also present in an intact copy. However, targeted analyses of particular genes of interest identified a few additional pseudogenes missed by Pseudofinder (Table S3; Fig. S10).

We used 45 conserved proteins encoded by the genome of *P. baltica* to analyse its relationship to other Rickettsiales members by a phylogenomic approach. Thanks to a substantially expanded sampling across the “Megaira group”, our phylogeny (Fig. 5) provided an important update compared to the previous analogous analysis by Davison et al. (2023). First, the endosymbiont of the ciliate *Hartmannula sinica* (“MegHsini1”), previously considered a potential member of the “Megaira group”, was robustly excluded from it and instead positioned sister to a lineage of Rickettsiaceae hosted by Nephromycea (a subgroup of Apicomplexa). Second, our analysis indicated that the “Megaira group” MAGs JAIELT01 and JAFLDA01 (*Ca*. Penulousia solacaria) represent bacteria conspecific with *Ca*. Megaira venefica and the endosymbiont of the ciliate *Ichthyophthirius multifiliis* strain G5, respectively. Third and most important, the “Megaira group” Clade B, not represented in the previous analysis, was recovered as monophyletic and fully supported. At the same time, the internal diversity of Clade B and the distinctiveness of *P. baltica* from other sequenced bacteria or available MAGs is clearly apparent from our tree (Fig. 5). The novelty of the *Ciliophrys* sp. Baltic endosymbiont is additionally underscored by the comparison of profiles of genes with KEGG annotations between sequenced “Megaira group” representatives (Table S4).

We analysed the gene repertoire of *P. baltica* to illuminate its biological features. A notable insight is that *P. baltica* lacks the ability to make flagella, unlike some other obligate intracellular bacteria, including some “Megaira group” representatives and other Rickettsiaceae (Davison et al. 2023; Castelli et al. 2024). On the other hand, unlike many members of the “Megaira group”, *P. baltica* encodes components of the protein machinery mediating the synthesis of O-antigen, i.e. a category of polymers of modified saccharides defining the surface characteristics of bacterial cell as the highly variable part of lipopolysaccharides. Many proteins encoded by the *P. baltica* genome do not have functionally characterized homologs, so their contribution to the biology of the bacterium cannot be assessed. Some do not even have any homologs discernible by simple BLASTp searches outside *P. baltica*, whereas others may exhibit unusual occurrence patterns, as exemplified by two hypothetical proteins (AC2SM2_00045 andAC2SM2_00050) with homology exclusively to proteins from *Phycorickettsia trachydisci* (a rickettsiacean endosymbiont of eustigmatophytes).

The set of metabolic pathways of *P. baltica*, as reconstructed by our genome analysis, is functionally reduced and generally reminiscent of the metabolic map of previously characterized Rickettsiaceae members (Fig. S11). Glycolysis is non-functional because of the absence of most involved enzymes, including pyruvate kinase. Of the gluconeogenesis pathway, only pyruvate, orthophosphate dikinase (PPDK) is present, apparently to produce phosphoenolpyruvate required for a particular step of lipopolysaccharide synthesis. The latter process also seems to be the main reason for *P. baltica* having preserved the non-oxidative part of the pentose phosphate pathway, as other common pathways that would rely on the provision of pentoses, including purine, pyrimidine, and histidine biosynthesis, are all absent from the bacterium. The majority of Rickettsiaceae rely on the supply of isoprenoid precursors from the host (Driscoll et al. 2017; Castelli et al. 2021), and the same holds for *P. baltica*, as it lacks the enzymes of both alternative pathways (MVA and MEP). The pathways for the synthesis of most proteinogenic amino acids, as well as thiamine, riboflavin, and biotin, are likewise missing. Because the host itelf is auxotrophic for many of the amino acids and all these cofactors, *P. baltica* must hijack these substances from the host’s diet.

The endosymbiont encodes all five NTP/NDP carrier proteins (Tlc1–Tlc5; Table S5), which enables the bacterium to import not only host-derived ATP in exchange for ADP, but also to transport other ribonucleotides (Audia & Winkler 2006). The energetic demands of the bacterium may, however, be primarily satisfied by metabolizing pyruvate via the standard route involving pyruvate dehydrogenase complex, the tricarboxylic acid (TCA) cycle, the respiratory chain, and ATP synthase (Fig. S11). *P. baltica* possesses all four conventional respiratory chain complexes, including cytochrome *c* oxidase (complex IV), while cytochrome *bd* ubiquinol oxidase, i.e. an alternative terminal oxidase, is apparently dysfunctional in this bacterium due to an in-frame termination codon in the subunit I gene (Table S3). Interestingly, a recent transcriptomic analysis revealed down-regulation of the core energetic metabolism of a ciliate host upon infection by the related bacterium *M. polyxenophila*, which the authors interpreted as such that the Krebs cycle and oxidative phosphorylation of the endosymbiont may provide ATP for both partners, with the NTP/NDP carriers functioning in the reverse than generally assumed (Giovannini et al. 2026). However, whether this apparent analogy to the energy metabolism integration of the mitochondrion with its “host” eukaryotic cell really holds for the *M. polyxenophila*-ciliate system or any other endosymbiotic partnership involving a rickettsialean bacterium such as *P. baltica*, needs to be corroborated. Unlike its relatives in the “Megaira group” or the host *Ciliophrys* sp. Baltic, *P. baltica* may, in principle, energetically benefit from histidine, as it possesses the histidine utilization (Hut) pathway. It comprises four enzymes that collectively convert histidine to formamide and L-glutamate, the latter of which is usable for proteosynthesis as well as metabolizable to pyruvate (Fig. 6; Table S6). All four enzymes constituting the Hut pathway are encoded by a single gene cluster (putative operon) that the *P. baltica* lineage most likely acquired by HGT from a gammaproteobacterial donor (Figs S12–S15). Since *Ciliophrys* sp. Baltic does not synthesize histidine *de novo, P. baltica* must obtain it indirectly through the host’s diet.

**Fig. 6.**
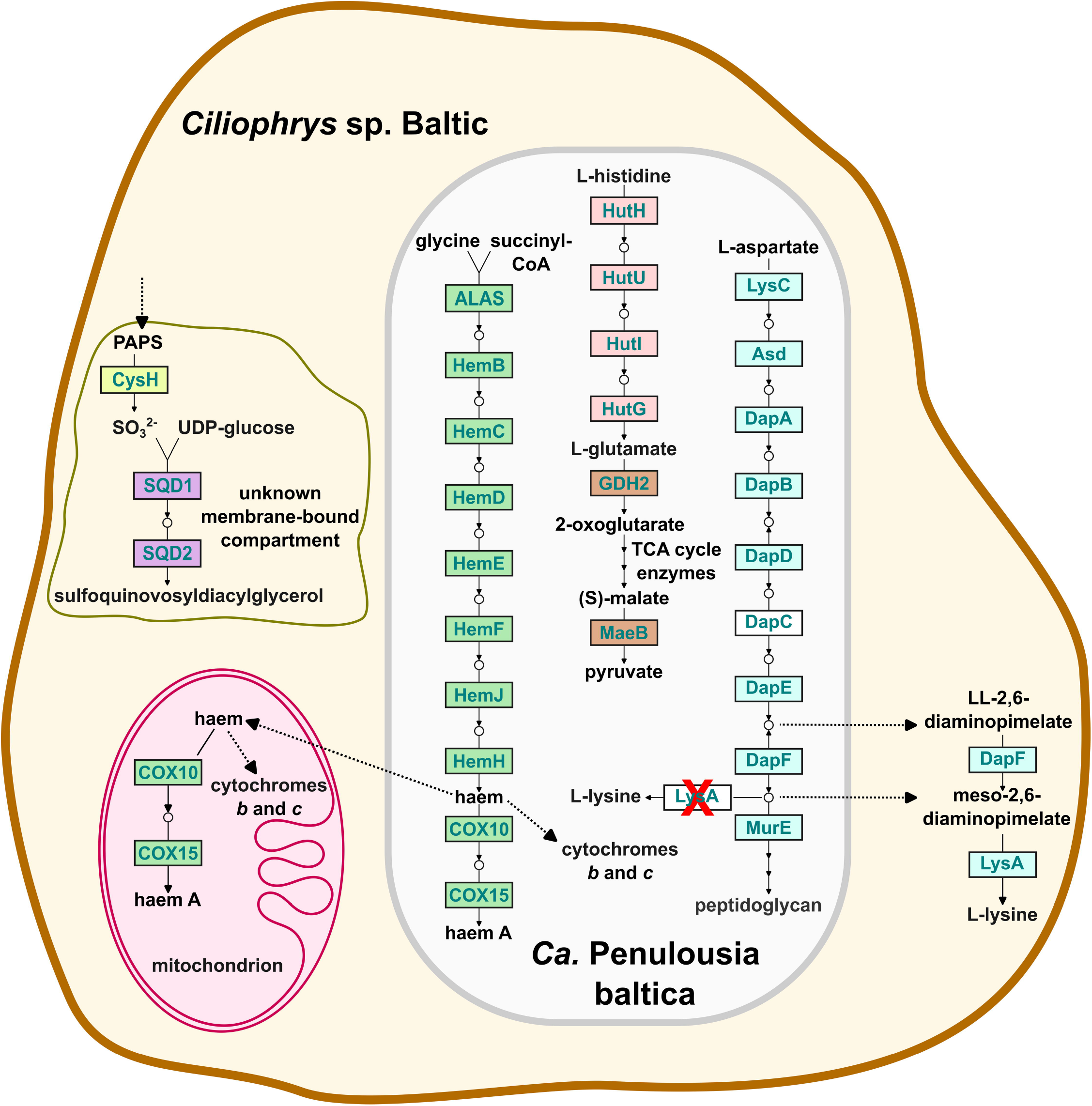
Metabolic pathways of interest in *Ca*. Penulousia baltica. The enzymes comprising the pathways are indicated with the respective designations as used in the KEGG database (for protein sequence IDs, see Tables S1 and S6). For simplicity, only the starting metabolites, the most relevant intermediates, and final products of the pathways are indicated in the scheme. The haem biosynthesis pathway is fully conserved in the bacterium to supply a cofactor needed to meet its own metabolic requirements. However, it may potentially also supply haem to the host *Ciliophrys* sp. Baltic, which is incapable of its synthesis. Diaminopimelate synthesised in *P. baltica* serves as one of the precursors of peptidoglycan but is not converted to lysine due to the lack of diaminopimelate decarboxylase (LysA crossed in red). The host cannot synthesize lysine *de novo* but may benefit from diaminopimelate leakage from the endosymbiont as it has retained the last two enzymes of the lysine biosynthesis pathway. Note the pathway hole, i.e., the absence of the enzyme N-succinyldiaminopimelate aminotransferase (DapC or ArgD), in the pathway leading to diaminopimelate in *P. baltica*, which is shared, e.g., with the genus *Rickettsia* and indicates the existence of an unknown alternative aminotransferase catalysing the reaction (Driscoll et al. 2017). The histidine utilization (Hut) pathway is unique for *P. baltica* among all other “Megaira group” members investigated and was acquired by an *P. baltica* ancestor via HGT from a gammaproteobacterial donor. The four enzymes of the Hut pathway endow the bacterium with the ability to convert histidine (putatively imported from the host via a transporter encoded by the same operon) to glutamate, which can then (among other uses) be metabolised to pyruvate for ATP production.

Two additional aspects of the reconstructed *P. baltica* metabolic map are particularly notable with regard to the metabolic integration of the endosymbiont and its host (Fig. 6; Table S6). First, the endosymbiont has retained a complete haem biosynthesis pathway, and thus synthesizes an essential compound for which its host is auxotrophic. We speculate that *Ciliophrys* sp. Baltic may have access to haem synthesized by *P. baltica*, potentially due to its release from endosymbionts following accidental disruption, active digestion of the endosymbionts by the host (perhaps via the autophagy pathway), or even its direct export from the endosymbiont via an unidentified transporter. Such a positive contribution to the host fitness by a Rickettsiales endosymbiont via providing an essential substance would not be unprecedented. For example, a strain of *Wolbachia* has been shown to provide B vitamins to its insect host, a function essential for host growth and reproduction (Nikoh et al. 2014; Newton & Rice 2020). An even closer relative of *P. baltica*, namely a *M. polyxenophila* strain occurring in the ciliate *Paramecium primaurelia*, was recently suggested to supply its host with biotin and explicitly designated as a facultative mutualist (Giovannini et al. 2026). We are unaware of a haem auxotroph other than *Ciliophrys* sp. Baltic that would host a rickettsialean endosymbiont, but it is notable that a betaproteobacterial endosymbionthosted by a lineage of parasitic trypanosomatids provides a biosynthetic haem precursor that is then converted to haem by the host, complementing the reactions missing in the endosymbiont (Kořený et al. 2013; Hammond et al. 2026). Such metabolic cooperation might also be in place in *P. baltica* and *Ciliophrys* sp. Baltic, securing the provision of lysine to both partners. Specifically, the bacterium would provide diaminopimelate, which it synthesizes as one of the precursors of peptidoglycan but apparently does not divert it for lysine production due to the lack of diaminopimelate decarboxylase (as is the case in Rickettsiaceae in general). On the other hand, the host has lost the *de novo* lysine biosynthesis pathway (together with the plastid) but still possesses enzymes for converting diaminopimelate to lysine (see above), so it can potentially benefit from the leakage of diaminopimelate from the endosymbiont (Fig. 6). An analogous metabolic cooperation to complete lysine biosynthesis has been observed between certain sap-feeding insects and their bacterial endosymbionts (Husnik et al. 2013; Luan et al. 2015).

### The Ca. Penulousia baltica genome encodes a diverse set of proteins potentially involved in host-symbiont interactions

The interaction between *P. baltica* and its host is not restricted to the metabolite exchange. Like other Rickettsiales (e.g., Gillespie et al. 2010), *P. baltica* harbours a type IV secretion system (T4SS), a versatile molecular apparatus commonly involved in the translocation of effector proteins or DNA into host cells (Fronzes et al. 2009). Specifically, the genome encodes the VirB/D4-like (P-type) components (Table S5). We additionally identified all three components of the HlyB-HlyD-TolC tripartite type I secretion system (T1SS; Table S5), indicating the presence of a distinct secretion machinery capable of exporting toxins or other effector proteins across the bacterial envelope (Bui et al. 2023). Furthermore, we detected an array of proteins with signal peptides (Tables S7 and S8) that are translocated outside the cell via the general Sec or Tat pathways, and eight proteins containing autotransporter domains (Table S7), which are known to mediate the secretion of virulence factors to the bacterial outer surface (Henderson & Nataro 2001). *P. baltica* also encodes a palatin-like phospholipase family protein (AC2SM2_00065), which promotes escape from host membranes and facilitates cytoplasmic colonization by preventing autophagic targeting (Borgo et al. 2022). Immediately downstream the corresponding gene, the bacterium also encodes all three components and additional accessory proteins of an alpha-pore-forming tripartite toxin MakABE, a highly disruptive agent primarily involved in actin filament disruption and Golgi fragmentation (Nadeem et al. 2021). No other sequenced Rickettsiales bacteria contain homologs of these proteins, suggesting a unique evolutionary adaptation of *P. baltica* to its host. The set of candidate effector proteins potentially secreted by *P. baltica* into its host also include proteins with the characteristic repeat motifs that fold into domains mediating protein–protein interactions and are often associated with host manipulation in endosymbiotic bacteria. Specifically, we identified at least 29 proteins containing ankyrin (ANK) repeats, five proteins containing leucine-rich repeats (LRRs), and 27 proteins with tetratricopeptide repeats (TPRs), and most of them were predicted to be extracellular using DeepLocPro (Table S5).

Also noteworthy are several type II toxin-antitoxin (TA) systems in the *P. baltica* genome. They are composed of a stable toxin and its cognate, typically unstable, antitoxin that neutralizes the toxin. We specifically identified both components of MazEF, VapBC, HipBA, and RelBE systems (Table S9). We further detected the RatA toxin of the type II RatAB TA, although the corresponding antitoxin could not be confirmed. It has been recently proposed that toxin-antitoxin systems encoded by endosymbiotic bacteria may addict the host to the endosymbionts (Midha et al. 2021; Castelli et al. 2024). In addition, the *P. baltica* genome encodes the type IV AbiE TA system, comprising the nucleotidyltransferase toxin AbiEii and its non-interacting antitoxin AbiEi (Table S9). This system likely functions to protect the endosymbiont from bacteriophage infections by inducing abortive infection in phage-infected cells, thereby preventing the spread of the virus within the population (Dy et al. 2014). We also identified a NACHT domain-containing protein (AC2SM2_00140), which is similarly known to defend against phages (Kibby et al. 2023). Furthermore, the genome of *P. baltica* contains so-called gene-transfer agents (GTAs), virus-like particles that mediate HGT. We identified all GTA components reported previously from other Rickettsiales (George et al. 2022), including the large terminase, portal and major capsid proteins, as well as tail proteins (Table S10). The role of GTAs in obligate endosymbionts remains unknown, but they could play a role in mediating interactions between the bacteria and their host.

Surprisingly, *P. baltica* contains a two-gene operon encoding a LodA/GoxA family CTQ-dependent oxidase (AC2SM2_00110) and a flavoprotein of the LoxB/GoxB type (AC2SM2_00115). The LodA/GoxA family enzymes use a protein-derived cofactor called CTQ (cysteine tryptophylquinone), which is generated by the action of the LodB/GoxB flavoprotein, to catalyze oxidative deamination reactions, typically converting primary amines (such as lysine or glycine) to aldehydes, with the concomitant release of ammonium and hydrogen peroxide (Lucas-Elío et al. 2006; Chacón-Verdú et al. 2014). The operon comprising genes homologous to *lodA*/*goxA* and *lodB*/*goxB* has been identified across diverse bacterial taxa including marine species, yet our survey of sequenced Rickettsiales members indicated its presence in *P. baltica* only. Horizontal transfer of the operon into the *P. baltica* lineage is therefore likely, although no specific donor lineage could be determined by phylogenetic analysis (Fig. S16). The LodA-like L-amino acid oxidases have been postulated to play roles in biofilm development and microbial competition, particularly through the production of H_2_O_2_ (Campillo-Brocal et al. 2015; Lucas-Elío et al. 2006). We cannot exclude the possibility that the *P. baltica* LodA/GoxA family protein is involved in biotic interactions of the bacterium, but it is noteworthy that it differs from the previously characterized lysine and glycine oxidases (LodA and GoxA, respectively) by possessing an N-terminal catalase-like domain. This architecture, to our knowledge, has not been described before, yet we found it in various other bacteria, e.g. the cyanobacterium *Acaryochloris marina* (WP_446077187.1). If the N-terminal domain has a *bona fide* catalase activity, it would perhaps detoxify H_2_O_2_ immediately upon its generation by the amine oxidase activity of the C-terminal part of the protein. Hence, the physiological role of biochemical activity of the LodA-like protein in *P. baltica* is not clear.

#### Perspectives

Endosymbiosis is a pervasive phenomenon and a major evolutionary factor of eukaryotic life. Our investigations of *Ciliophrys* sp. Baltic further highlight the different evolutionary trajectories that endosymbionts may follow. One category comprises obligate endosymbiosis characterized by a tight integration of the partners, vertical inheritance of the endosymbiont, and its progressive genome reduction (Husnik et al. 2021). The usual fate of such endosymbionts perhaps is their degradation and eventual loss (Husnik and Keeling 2019), with two prominent cases in this category being exceptional: mitochondria and plastids that have managed to persist for more than a billion years. Still, even in these cases specific circumstances may open a road to a complete endosymbiont (organelle) loss (Maciszewski and Karnkowska 2019). For plastids, such a critical step may be the loss of photosynthesis, although it is notable that of the documented cases of complete plastid loss only those concerning syndineans and the plastid-lacking apicomplexan taxa directly support this notion, as they are phylogenetically nested among plastid-bearing non-photosynthetic clades (Holt et al. 2023; Mathur et al. 2023). In contrast, the three other lineages previously documented to secondarily lack a plastid, i.e. Picozoa and the ochrophytes *P. flagellatus* and *A. sol*, do not have known non-photosynthetic sister groups, that is, there is no direct evidence for a stepwise plastid reduction with a non-photosynthetic plastid as a transition stage. Nevertheless, a direct loss of a photosynthetic plastid is an unlikely scenario, so it is possible that the expected “missing links” are eventually found. Herein, we demonstrated that the plastid has also vanished in a third ochrophyte lineage represented by *Ciliophrys* sp. Baltic. Its closest relatives whose plastid status is clearly defined are the photosynthetic Rhizochromulinaceae members, but we posit that a non-photosynthetic reduced plastid may still persist in representatives of Ciliophryaceae other than *Ciliophrys* sp. Baltic itself; work is in progress to test this possibility.

A salient question also is how much *Ciliophrys* sp. Baltic is representative of its whole family when it comes to the manifestation of the other major category of endosymbiosis, the one realized by the professional endosymbionts. Our analysis of the genome of the bacterium inhabiting *Ciliophrys* sp. Baltic, herein described as *Candidatus* Penulousia baltica, robustly documents its attribution among professional endosymbionts. While their relationship to the host is generally less beneficial compared to the obligate endosymbionts, being parasitic or commensalistic at best, our analyses pointed to the possibility that *Ca*. Penulousia baltica might confer a benefit to its host by providing haem or the lysine precursor diaminopimelate, which *Ciliophrys* sp. Baltic cannot synthesize itself due to the plastid loss. By obtaining and characterizing *Ciliophrys* sp. Baltic, we established a unique model system for studying functional and evolutionary aspects of endosymbiosis in protists.

### Description of Candidatus Penulousia baltica sp. nov

Candidatus Penulousia baltica (L. fem. adj. *baltica*, pertaining to the Baltic Sea). Endosymbiont of *Ciliophrys* sp. Baltic. The species is phylogenetically distinct from other described species in the genus. The type genome for the species is the GenBank assembly JBXYML000000000.1. The GC content of the type genome is 35.38% and the genome size is 1,513,035 bp.

## Materials and Methods

### Culturing, imaging, and sequencing

*Ciliophrys* sp. Baltic was isolated from a sand sample collected on August 18, 2022, at Melnragė Beach in Klaipėda, Lithuania (55.7376° N, 21.0856° E). The established strain was maintained in CCALA brackish water (https://ccala.butbn.cas.cz/media-receipts), with cereal grass enrichment used initially but later omitted, as the strain grew well without it. The cultures were kept at room temperature in filter-capped cell culture flasks under a natural day-night cycle and subcultured once per several months. Light micrographs were taken with an Olympus BX53 microscope equipped with a DP73 digital camera. Samples for fluorescence in situ hybridization (FISH) using the oligonucleotide probe CilioEndo16S (5’-GCGTCAGTTGAAGTCCAGGTG-3’), labelled with a 5’-Cy3 fluorescent dye, were prepared as detailed in George et al. (2023) and examined using the Olympus BX53 microscope with a U-RFL-T fluorescence light source. The TEM was performed as described in Barcytė et al. (2024). Total genomic DNA and RNA from *Ciliophrys* sp. Baltic were independently isolated using MasterPure Complete DNA & RNA Purification Kit (Biosearch Technologies) following the manufacturer’s instructions. The extracted DNA was initially subjected to both short-read Illumina NovaSeq 6000 (library preparation using the NEBNext Ultra II DNA Kit) and long-read Oxford Nanopore MinION (1D ligation kit) sequencing at the Sequencing Section of the Okinawa Institute of Science and Technology. RNA was also sequenced using the same Illumina platform, with a library prepared using the NEBNext Ultra II RNA Library Preparation kit. To increase sequencing coverage, newly extracted DNA and RNA of *Ciliophrys* sp. Baltic were sequenced using the NovaSeq X platform, with DNA processed by Eurofins Genomics (Constance, Germany) and RNA by Macrogen Europe (Amsterdam, Netherlands). For RNA, TruSeq Stranded mRNA LT Sample Prep Kit was used.

### Genome and transcriptome assemblies

Paired-end Illumina genomic reads were first trimmed and filtered using Trimmomatic v0.39 (Bolger et al. 2014) with the parameters −phred33 ILLUMINACLIP:2:20:10 LEADING:3 TRAILING:3 SLIDINGWINDOW:4:15 MINLEN:75. The post-processed reads were then co-assembled with Nanopore reads using SPAdes v3.15.5 (Bankevich et al. 2012). The latter were also assembled separately with Flye v2.9.1-b1780 (Kolmogorov et al. 2020) using -- meta argument. Inspection of the assembly graph with Bandage v0_8_1 (Wick et al. 2015), uncovered a cluster of contigs sharing similar coverage levels and totalling about 1.5 Mb. By performing BLASTn search (Altschul et al. 1990) of our Flye assembly using *Candidatus* Megaera polyxenophila genome (CP104166.1) as a query, we identified these contigs to belong to a Rickettsiales bacterium. We mapped the corresponding Nanopore reads to these contigs using minimap2 v2.17-r941 (Li 2018) and extracted them using samtools v1.15.1 (Li et al. 2009). Illumina reads corresponding to the Rickettsiales bacterium were retrieved using BBSplit (BBMap/BBTools package) using the identified contigs as well as CP104166.1 as references. All extracted Illumina and Nanopore reads were then co-assembled with Unicycler v0.4.8 using default settings (Wick et al. 2017). The newly obtained RNA sequencing reads were quality-trimmed in the same manner as the DNA sequencing reads and were then assembled with rnaSPAdes v3.15.5 (Bushmanova et al. 2019) using default parameters. TransDecoder v5.7.1 (https://github.com/TransDecoder/) with the LongOrfs utility and default settings was used to infer encoded protein sequences from the transcriptome assembly.

### Identification, annotation, and sequence analysis of Ciliophrys sp. Baltic genes

The initial annotation of the assembled mitogenome was obtained with MFannot (Lang et al. 2023) and subsequently refined through manual curation, including HHpred searches (Zimmermann et al. 2018) employed for a more sensitive homology detection where BLAST failed to retrieve homologous sequences. The tertiary structure of the protein encoded by the mitochondrial gene *orf211* was modelled using AlphaFold 3 (https://alphafoldserver.com/; Abramson et al. 2024) and ESMFold (https://www.tamarind.bio/tools/esmfold; Lin et al. 2023) and the resulting models were used as queries for FoldSeek searches (https://search.foldseek.com/; van Kempen et al. 2024). A graphical map of the mitogenome was generated with OGDRAW v1.3.1 (Greiner et al. 2019). The level of completeness of the *Ciliophrys* sp. Baltic transcriptome assembly and the closed *Ca*. P. baltica genome sequence was assessed using BUSCO v5.2.2 (Manni et al. 2021). Nuclear genes of interest from *Ciliophrys* sp. Baltic were searched in the transcriptome assembly and the full genome assembly using tBLASTn and appropriate reference sequences as queries. Specifically, we relied on previously characterized sequences from other ochrophytes (Barcytė et al. 2024; Terpis et al. 2025), on well-annotated genes from the diatom *Phaeodactylum tricornutum* as presented in the KEGG database (https://www.kegg.jp/kegg-bin/show_organism?menu_type=pathway_maps&org=pti), and on protein sequences inferred from available transcriptome assemblies from the closest *Ciliophrys* relatives (i.e., other dictyochophytes) to maximize the chance that the sought-after homologs were detected. Candidate homologs were checked by BLASTing them again the NCBI non-redundant (nr) protein sequence database or the KEGG database (https://www.genome.jp/tools/blast/) to filter out sequences derived from the organisms co-existing with *Ciliophrys* sp. Baltic (the endosymbiont and the co-cultured prokaryotes) and to verify the identification of sequences as orthologs of the query reference proteins. We primarily report sequences derived from the *Ciliophrys* sp. Baltic transcriptome assembly, but in the cases of the transcript contigs being truncated or unspliced, we reconstructed the full coding nucleotide sequence and the encoded amino acid sequence from the genome assembly, deducing the exon-intron structure based on comparison with homologous sequences (typically the respective dictyochophyte orthologs). To minimize the possibility that some of the hallmark plastid proteins are in fact present in *Ciliophrys* sp. Baltic yet are too divergent to be identified by BLAST, we employed the more sensitive HMMER 3 searches (Eddy 2011) against the set of protein sequences inferred from the transcriptome assembly. As queries we used profile HMMs derived from seed alignments of the respective records in the Pfam database, except for the Omp85 protein, for which we used a custom profile HMM built from a multiple alignment of selected ochrophyte representatives (see Figshare, doi: 10.6084/m9.figshare.32287122) to cope for the fact that this plastid protein is very divergent and is not generally found when the broadly defined Pfam family (dominated by bacterial Omp85 proteins) is employed. TargetP-2.0 (https://services.healthtech.dtu.dk/services/TargetP-2.0/; Almagro Armenteros et al. 2019) was employed for investigating the presence of different types of localization signals (signal peptide, mitochondrial transit peptide) in proteins of interest from *Ciliophrys* sp. Baltic and other dictyochophytes; the analyses were performed in the “non-plant” search mode and with default thresholds. To get a more specific prediction of the protein subcellular localization (see Table S5), we additionally used DeepLoc 2.1 (Ødum et al. 2024).

### Annotation and analysis of the Candidatus Penulousia baltica genome

The genome of the bacterial endosymbiont *Ca*. P. baltica was initially annotated using Prokka v1.12 (Seemann 2014) and visualized with Proksee (https://proksee.ca; Grant et al. 2023). Genes and proteins of interest from the *Ca*. P. baltica genome were searched in the genome sequence or the set of annotated proteins sequences using BLAST. Identified candidates were evaluated by BLAST searches against the NCBI nr database and the KEGG database to confirm their identity. The taxonomic composition of the bacterial community co-cultured with *Ciliophrys* sp. Baltic was assessed with phyloFlash (Gruber-Vodicka et al. 2020). Candidate pseudogenes in the bacterial genome were identified with Pseudofinder v1.1.0 (Syberg-Olsen et al. 2022). Manual curation refined the predicted set by comparing candidates with homologs from other taxa, confirming some as true pseudogenes while excluding most others. KEGG Orthology assignments were performed using blastKOALA (Kanehisa et al. 2016) and Anvi’o 7 (Eren et al. 2021) to characterize individual gene functions and reconstruct KEGG pathways. Pathway displays were generated with KEGG Mapper (Kanehisa et al. 2021). HMMER searches were used to detect proteins with domains composed of ankyrin, leucine-rich, and pentatricopeptide repeats. Toxin-antitoxin systems were identified with TAfinder 2.0 (Guan et al. 2024) and by BLAST searches. The repertoire of antiphage defense systems in *Ca*. P. baltica was assessed using DefenseFinder (https://defensefinder.mdmlab.fr/; Tesson et al. 2024), with the AbiE TA system being the only identified. Proteins with signal peptides were identified with SignalP 6.0 (Table S7; Teufel et al. 2022) and Phobius (Table S8; Käll et al. 2004), and the protein localization was additionally predicted using DeepLocPro (Moreno et al. 2024). While all the analyses reported in this study were carried out based on our annotation of the *Ca*. P. baltica genome, for the purpose of presenting the results (in the text, figure and tables) we use gene IDs derived from a parallel annotation generated by GenBank upon submission of the genome sequence to the databae. Our original annotation is provided in Figshare, doi: 10.6084/m9.figshare.32287122.

### Phylogenetic analyses

In general, phylogenetic analyses were performed on multiple sequence alignments created using MAFFT v7 (Katoh & Standley 2013) and trimmed to remove poorly aligned regions (details specified below for each analysis), which were then subjected to a maximum likelihood tree search implemented in IQ-TREE multicore v2.2.5 (Minh et al., 2020) with the Model Finder Plus setting (MFP; Kalyaanamoorthy et al. 2017) to select the best-fitting substation model (according to the Bayesian Information Criterion). The length of the final alignment, the selected substitution model, and the method of branch support evaluation (non-parametric or ultrafast bootstrapping) is indicated specifically for each tree in the respective figure legend. The phylogenetic position of the newly isolated *Ciliophrys* sp. Baltic strain was determined using the 18S rDNA sequence based on a dataset compiled to sample sequences from all major ochrophyte lineages, including all unique *Ciliophrys* genotypes represented in the GenBank nucleotide sequence database. One additional sequence was identified by exploring the Tara Oceans DADA2 ASVs 18S V4 (eukaryotes) database implemented in OBA. The 73 sequences were aligned with the L-INS-i algorithm and the alignment was trimmed with ClipKIT v2.6.1 in the gappy mode (Steenwyk et al. 2020). The phylogenetic analysis of the 16S rDNA sequences to determine the position of the endosymbiont (*Ca*. P. baltica) was carried out on a sequence dataset compiled by including closest matches in BLAST searches of relevant NCBI sequence databases (the nr/nt database and the wgs database restricted to sequences from Alphaproteobacteria), along with more dissimilar sequences selected based on previously published phylogenetic analyses to obtain a representative sample across the diversity of Rickettsiales. A sequence from a “Megaira group” representative accidentally identified in the previously reported transcriptome assembly from *Azumiobodo hoyamushi* (Yazaki et al. 2017) was included, too. The sequences were aligned using the E-INS-i MAFFT algorithm and the alignment was automatically trimmed with ClipKIT v2.6.1 in the gappy mode. To construct a phylogenomic dataset for Ricketsiales, we first used OrthoFinder (Emms & Kelly 2019) to identify single-copy orthologs across a broad taxonomic sampling of the group. This dataset included previously analysed complete or draft genome assemblies from known hosts and metagenome-assembled genomes (MAGs) from mixed environmental samples (Hunter et al. 2020; Davison et al. 2023; Wittmers et al. 2025) further supplemented by additional selected rickettsialean genome assemblies and MAGs identified by tBLASTn searches in the nr/nt and wgs NCBI databases as putatively being related to the *Ciliophrys* sp. Baltic endosymbiont (see the list in Table S11). Furthermore, as we found that a full genome assembly of the brown alga *Ectocarpus* sp. 12 AP-2014 (CAXNAL020000000) contains sequences apparently coming from an endosymbiont related to the bacterium in *Ciliophrys* sp. Baltic, we included into the analysis also the set of protein sequences derived from the full assembly (assuming that the sequences from the rickettsialean endosymbiont will be fished out using our procedure). Finally, sequences from an unpublished genome assembly of the endosymbiont of *Ichthyophthirius multifiliiis* strain G5 (see Sun et al. 2009) kindly provided to us by Dr. Thomas Doak (Department of Biology, Indiana University, Bloomington), were included, too. Due to the low number of single-copy orthologs initially recovered, we manually selected additional genes, resulting in a final set of 45 orthologs. The sets of orthologous sequences were aligned separately with the E-INS-i method. The alignments were trimmed with trimAl v1.2rev59 using the -gappyout option (Capella-Gutiérrez et al. 2009) and concatenated into a single supermatrix using FASconCAT-G v1.05 (Kück & Longo 2014). For derlin protein (Der1-1 and Der1-2) phylogeny, the original dataset of Ponce-Toledo et al. (2025) was enriched with sequences from *Ciliophrys* sp. Baltic and additional representative non-photosynthetic ochrophytes (*Paraphysomonas imperforata, Leucomyxa plasmidifera, Picophagus flagellatus*, and *Actinophrys sol*) identified by BLAST searches (for the sequence IDs and source datasets see Table S12). All sequences were aligned with the E-INS-i algorithm and trimmed with ClipKIT in the gappy mode. Alignments of Hsp70, FabD, and FabI proteins were prepared in the same way using datasets of selected closest homologs identified by BLAST searches against NCBI. For the Hut operon (hutG, hutH, hutI, and hutU), each gene was queried against NCBI using BLAST, and the 100 best hits were downloaded for phylogenetic analyses. For hutG and hutU, one and three additional sequences, respectively, were included. Alignments for these single-gene phylogenies were prepared using the same method as for the derlin proteins. For the LodA/GoxA phylogeny, an additional 285 homologs were collected using BLAST and processed as described above for the other genes.

## Data availability

Sequencing reads from the *Ciliophrys* sp. Baltic culture were deposited at the National Center for Biotechnology Information as the BioProject PRJNA1460246. The 18S rDNA and mitogenome sequences of *Ciliophrys* sp. Baltic, as well as the annotated *Ca*. Penulousia baltica genome sequence, were deposited in GenBank under accession numbers PZ341965.1, PZ337301.1, and JBXYML000000000.1, respectively. Genome and transcriptome assemblies from the *Ciliophrys* sp. Baltic culture (containing sequences of the eukaryotic host, its endosymbiont, and co-cultured other bacteria), together with sequence alignments, are available from Figshare doi: 10.6084/m9.figshare.32287122.

## Ethics declarations

The authors declare no competing interests.

## Contributions

Conceptualization: D.B., M.E.; Data curation: D.B.; Formal analysis: D.B., M.E.; Funding acquisition: M.E., F.H.; Investigation: D.B.; Writing – original draft: D.B., M.E.; Writing – review & editing: F.H.

## Acknowledgements

The work was supported by the Czech Science Foundation project 23-06203S (to ME). DB and ME were also supported by the European Union under the LERCO project number CZ.10.03.01/00/22_003/0000003 via the Operational Programme Just Transition. FH is partly supported by an HFSP Early Career Grant (RGEC29/2024; https://doi.org/10.52044/HFSP.RGEC292024.pc.gr.194160). We thank the Scientific Computing (SCDA) and Sequencing (SQC) sections of the core facilities at the Okinawa Institute of Science and Technology for their great support.

## Supplementary figures

**Fig. S1**. Full version of the ML phylogenetic tree inferred from nuclear 18S rRNA gene sequences. Further details can be found in the legend to Fig. 1, which shows a simplified version of the tree.

**Fig. S2**. ML phylogenetic tree inferred from derlin protein (Der1-1 and Der1-2) sequences. The original dataset of Ponce-Toledo et al. (2025) was enriched with additional sequences (marked in bold). The final alignment consisted of 455 sequences with 343 aligned positions. The phylogenetic tree was inferred with IQ-TREE using LG+C20+G+I substitution model. Numbers at branches indicate bootstrap support values (>50%) calculated from 1,000 ultrafast bootstrap replicates. Der1-1 and Der1-2 paralogs are highlighted in rose and green colours, respectively. SELMA sequences are boxed in a more saturated colour.

**Fig. S3**. ML phylogenetic tree of nucleus-encoded Hsp70 isoforms. The final alignment consisted of 152 sequences and 774 aligned positions. The phylogenetic tree was inferred using IQ-TREE with the Q.YEAST+F+I+R7 substitution model. Numbers at branches represent bootstrap values (shown when >50%) calculated from 100 non-parametric bootstrap replicates. Homologs from *Ciliophrys* sp. Baltic, for which sequence IDs correspond to genome scaffolds, are shown in bold.

**Fig. S4**. Maximum-likelihood phylogenetic tree of FabD proteins. The final alignment comprised 88 sequences and 406 aligned positions. The phylogenetic tree was inferred using IQ-TREE under the Q.PFAM+R6 substitution model. Numbers at branches indicate bootstrap support values (>50%) calculated from 100 non-parametric bootstrap replicates. The sequence from *Ciliophrys* sp. Baltic is shown in bold (the sequence ID is provided in Table S1). Plastid homologs from other Dictyochophyceae are highlighted with a light pink box.

**Fig. S5**. Maximum-likelihood phylogenetic tree of FabI proteins. The final alignment comprised 75 sequences and 371 aligned positions. The phylogenetic tree was inferred using IQ-TREE under the Q.PFAM+R6 substitution model. Numbers at branches indicate bootstrap support values (>50%) calculated from 100 non-parametric bootstrap replicates. The sequence from *Ciliophrys* sp. Baltic is shown in bold (the sequence ID is provided in Table S1). Plastid homologs from other Dictyochophyceae are highlighted with a light pink box.

**Fig. S6**. N-terminal sequences of selected plastid-targeted enzymes from dictyochophytes, including their orthologs from *Ciliophrys* sp. Baltic. (A) FabD; (B) FabI; (C) DapF; (D) LysA. Amino acid residues corresponding to N-terminal presequences, namely the signal peptide (in plastid-targeted proteins) and the mitochondrial transfer peptide (in mitochondrion-targeted proteins) as predicted by TargetP-2.0 are highlighted in violet and orange, respectively. The residues immediately following the presequences (i.e., the N-termini exposed upon their cleavage) are boxed in red. Note the absence of any N-terminal presequences in the DapF and LysA proteins from *Ciliophrys* sp. Baltic. Furthermore, as in *Ciliophrys* sp. Baltic, the ortholog of the plastid-targeted FabD in the non-photosynthetic species *Pteridomonas danica* is also predicted to possess a mitochondrial transfer peptide, indicating an independent remodelling of the N-terminal region and a rerouting of the protein to the mitochondrion.

**Fig. S7**. N-terminal sequences of SQDG biosynthesis enzymes. Residues that are immediately downstream of the predicted cleavage site of signal peptide processing peptidase (which removes the N-terminal signal peptide in the ER lumen) are boxed in red.

**Fig. S8**. phyloFlash output demonstrating the presence of a variety of bacteria in the (meta)genome data from *Ciliophrys* sp. Baltic.

**Fig. S9**. Full version of the ML phylogenetic tree inferred from Rickettsiales 16S rRNA gene sequences. Further details can be found in the legend to Fig. 4, which shows a simplified version of the tree.

**Fig. S10**. Genome map of *Ca*. Penulousia baltica. The outermost circle shows the positions and orientations of protein-coding genes (CDSs). The second circle indicates tRNA genes, and the third circle shows putative pseudogenes. The fourth and the fifth indicate rRNA and tmRNA genes, respectively. The innermost circles display the GC content (black line) and GC skew (green/purple line), highlighting regions of compositional bias.

**Fig. S11**. An overview of selected metabolic pathways of *Ca*. Penulousia baltica generated with KEGG Mapper. The schemes indicate the occurrence of particular enzymes in the bacterium (the rectangles in green) as reconstructed using a BlastKOALA search against the KEGG database.

**Fig. S12**. Maximum-likelihood phylogenetic tree of hutH homologs. The final alignment included 101 sequences and 514 positions. The substitution model employed was LG+I+G4 as automatically selected by IQ-TREE. Numbers at branches represent bootstrap values (shown when >50%) calculated from 100 non-parametric bootstrap replicates.

**Fig. S13**. Maximum-likelihood phylogenetic tree of hutU homologs. The final alignment included 104 sequences and 565 positions. The substitution model employed was LG+I+G4 as automatically selected by IQ-TREE. Numbers at branches represent bootstrap values (shown when >50%) calculated from 100 non-parametric bootstrap replicates.

**Fig. S14**. Maximum-likelihood phylogenetic tree of hutI homologs. The final alignment included 101 sequences and 425 positions. The substitution model employed was Q.pfam+I+G4 as automatically selected by IQ-TREE. Numbers at branches represent bootstrap values (shown when >50%) calculated from 100 non-parametric bootstrap replicates.

**Fig. S15**. Maximum-likelihood phylogenetic tree of hutG homologs. The final alignment included 102 sequences and 341 positions. The substitution model employed was LG+F+I+G4 as automatically selected by IQ-TREE. Numbers at branches represent bootstrap values (shown when >50%) calculated from 100 non-parametric bootstrap replicates.

**Fig. S16**. Phylogenetic analysis of LodA/GoxA homologs. The final alignment included 286 sequences and 1042 positions. The substitution model employed was Q.pfam+R10 as automatically selected by IQ-TREE. Numbers at branches represent bootstrap values (shown when >50%) calculated from 10,000 ultrafast bootstrap replicates. The resulting phylogeny did not clearly resolve the donor lineage of the LodA/GoxA gene horizontally transferred into the *Candidatus* Penulousia baltica genome.

## Supplementary Tables

**Table S1**. Nuclear-genome encoded proteins from *Ciliophrys* sp. Baltic discussed in this study.

**Table S2**. Hallmark nucleus-encoded plastid proteins in selected ochrophytes.

**Table S3**. Putative pseudogenes detected in *Candidatus* Penulousia baltica genome.

**Table S4**. Comparison of KEGG-annotated gene profiles among sequenced representatives of the “Megaira group”, highlighting the distinct functional repertoire of *Ca*. Penulousia baltica (the Ciliophrys sp. Baltic endosymbiont). The original table published by Davison et al. (2023) was updated by including *Ca*. Penulousia baltica, for which KEGG annotations were generated using two independent approaches, BlastKOALA and Anvi’o, and are presented in separate columns. Highlights indicate disagreements between the two annotation approaches.

**Table S5**. Proteins putatively associated with host manipulation and virulence detected in the *Ca*. Penulousia baltica genome.

**Table S6**. Enzymes of metabolic pathways of special interest in *Ca*. Penulousia baltica.

**Table S7**. Predicted signal peptides in the *Ca*. Penulousia baltica proteome identified using SignalP 6.0. The table lists proteins predicted to contain signal peptides, their predicted type, and associated prediction scores. The identity of each protein based on BLAST searches against NCBI is also provided, along with the corresponding protein sequence.

**Table S8**. Predicted signal peptides in the *Ca*. Penulousia baltica proteome identified using Phobius. The table lists proteins predicted to contain signal peptides along with associated prediction scores. For proteins not predicted by SignalP 6.0 (Table S7), the protein identity and corresponding sequence are also provided.

**Table S9**. Toxin-antitoxin systems identified in *Ca*. Penulousia baltica genome.

**Table S10**. Gene-transfer agent (GTA) genes identified in the genome of *Ca*. Penulousia baltica.

**Table S11**. Bacterial genomes and MAGs included in the phylogenomic analysis.

**Table S12**. Sequence IDs and source information for derlin homologs added to the previously published dataset (Ponce-Toledo et al. 2025) for the purpose of phylogenetic analysis.

