## Supplementary figures for "The colourless dictyochophyte *Ciliophrys* sp. Baltic secondarily lacks a plastid but hosts a bacterial endosymbiont"

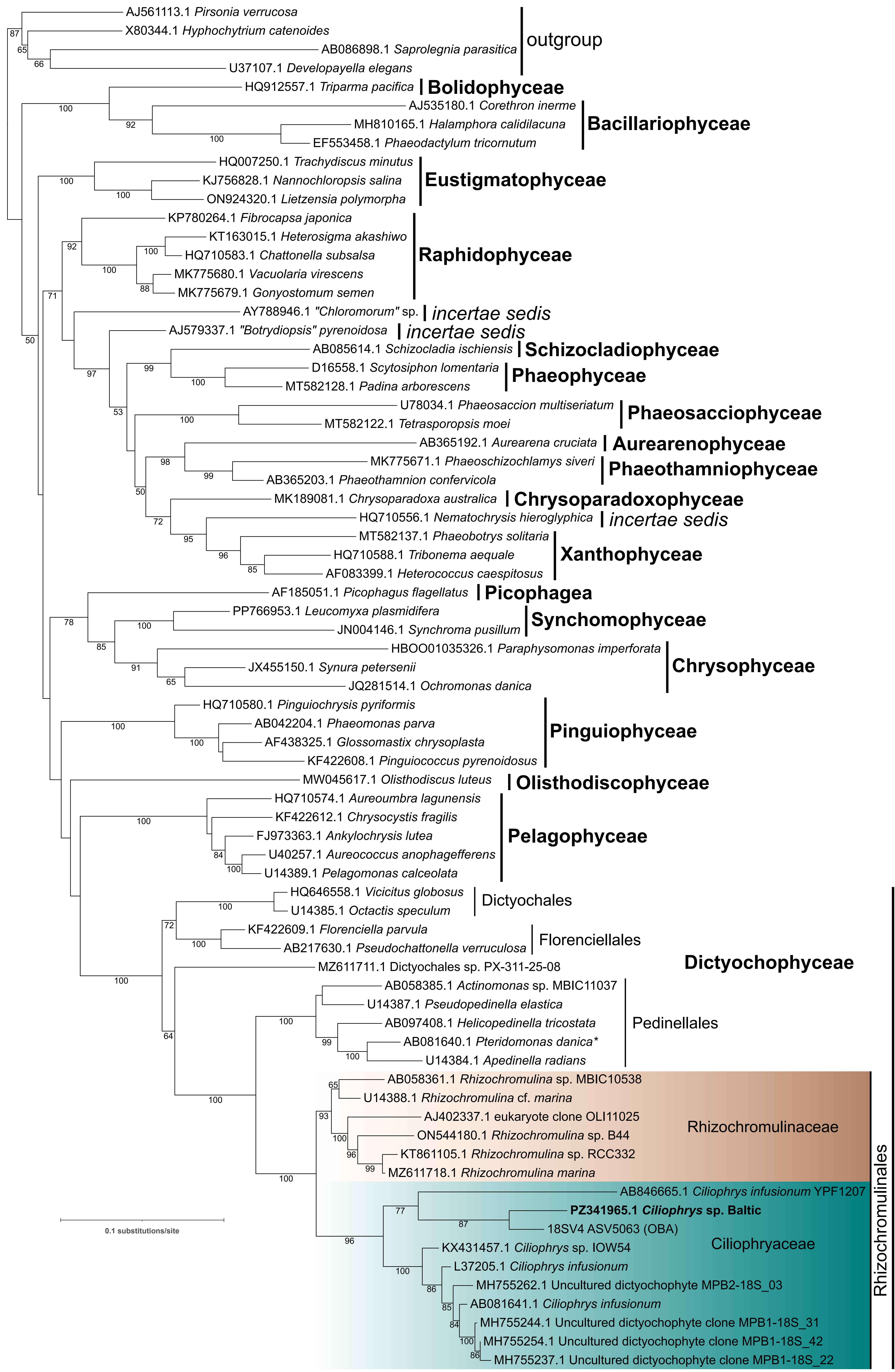


**Fig. S1.** Full version of the ML phylogenetic tree inferred from nuclear 18S rRNA gene sequences. Further details can be found in the legend to Fig. 1, which shows a simplified version of the tree.

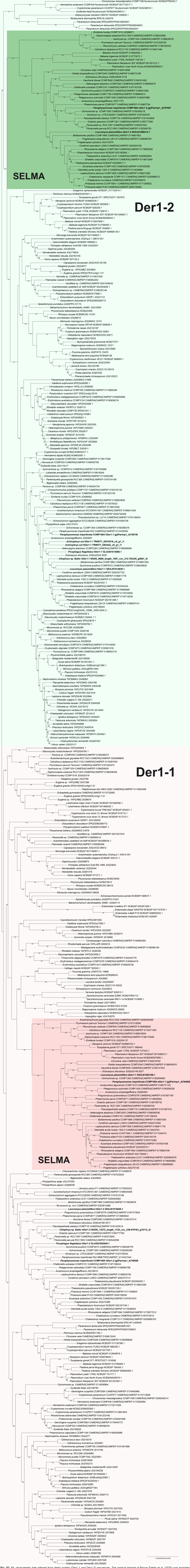


**Fig. S2.** ML phylogenetic tree inferred from derlin protein (Der1-1 and Der1-2) sequences. The original dataset of Ponce-Toledo et al. (2025) was enriched with additional sequences (marked in bold). The final alignment consisted of 455 sequences with 343 aligned positions. The phylogenetic tree was inferred with IQ-TREE using LG+C20+G+I substitution model. Numbers at branches indicate bootstrap support values (>50%) calculated from 1,000 ultrafast bootstrap replicates. Der1-1 and Der1-2 paralogs are highlighted in rose and green colours, respectively. SELMA sequences are boxed in a more saturated colour.

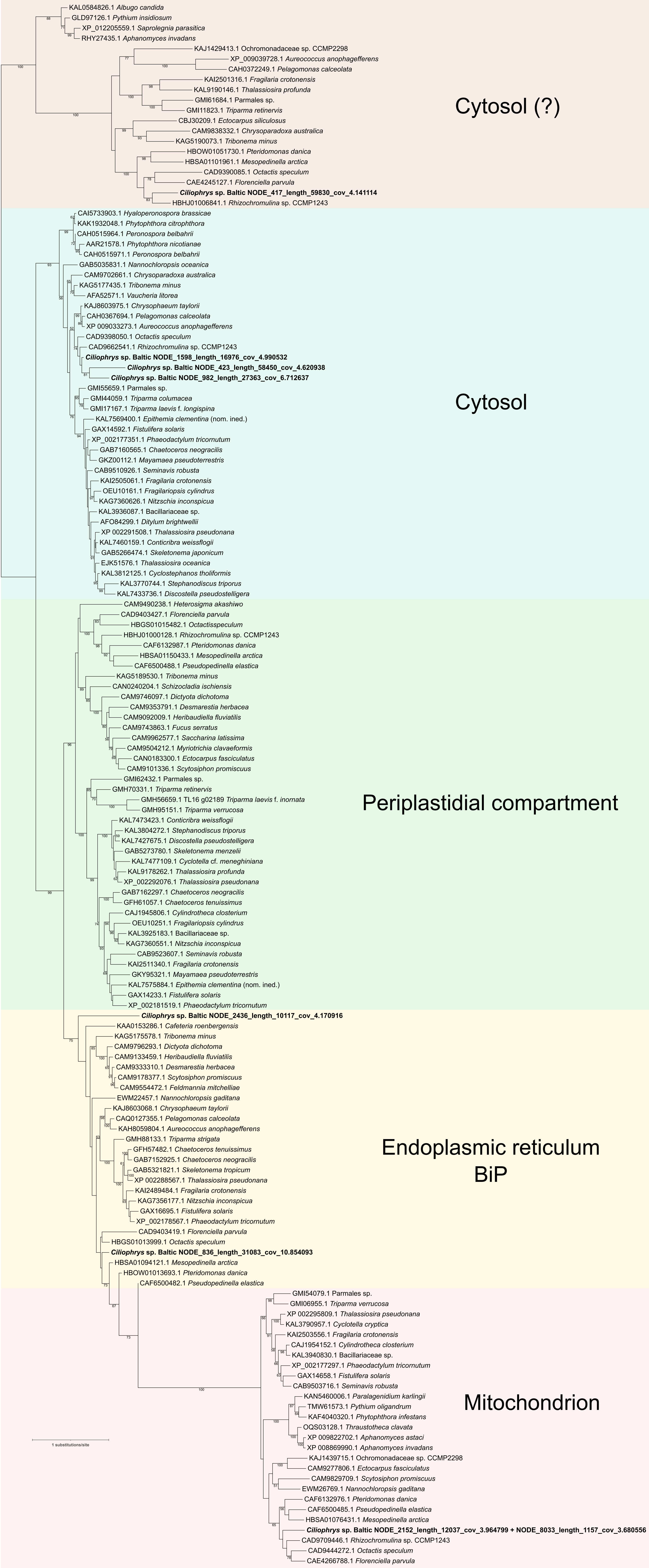


**Fig. S3.** ML phylogenetic tree of nucleus-encoded Hsp70 isoforms. The final alignment consisted of 152 sequences and 774 aligned positions. The phylogenetic tree was inferred using IQ-TREE with the Q.YEAST+F+I+R7 substitution model. Numbers at branches represent bootstrap values (shown when >50%) calculated from 100 non-parametric bootstrap replicates. Homologs from *Ciliophrys* sp. Baltic, for which sequence IDs correspond to genome scaffolds, are shown in bold.

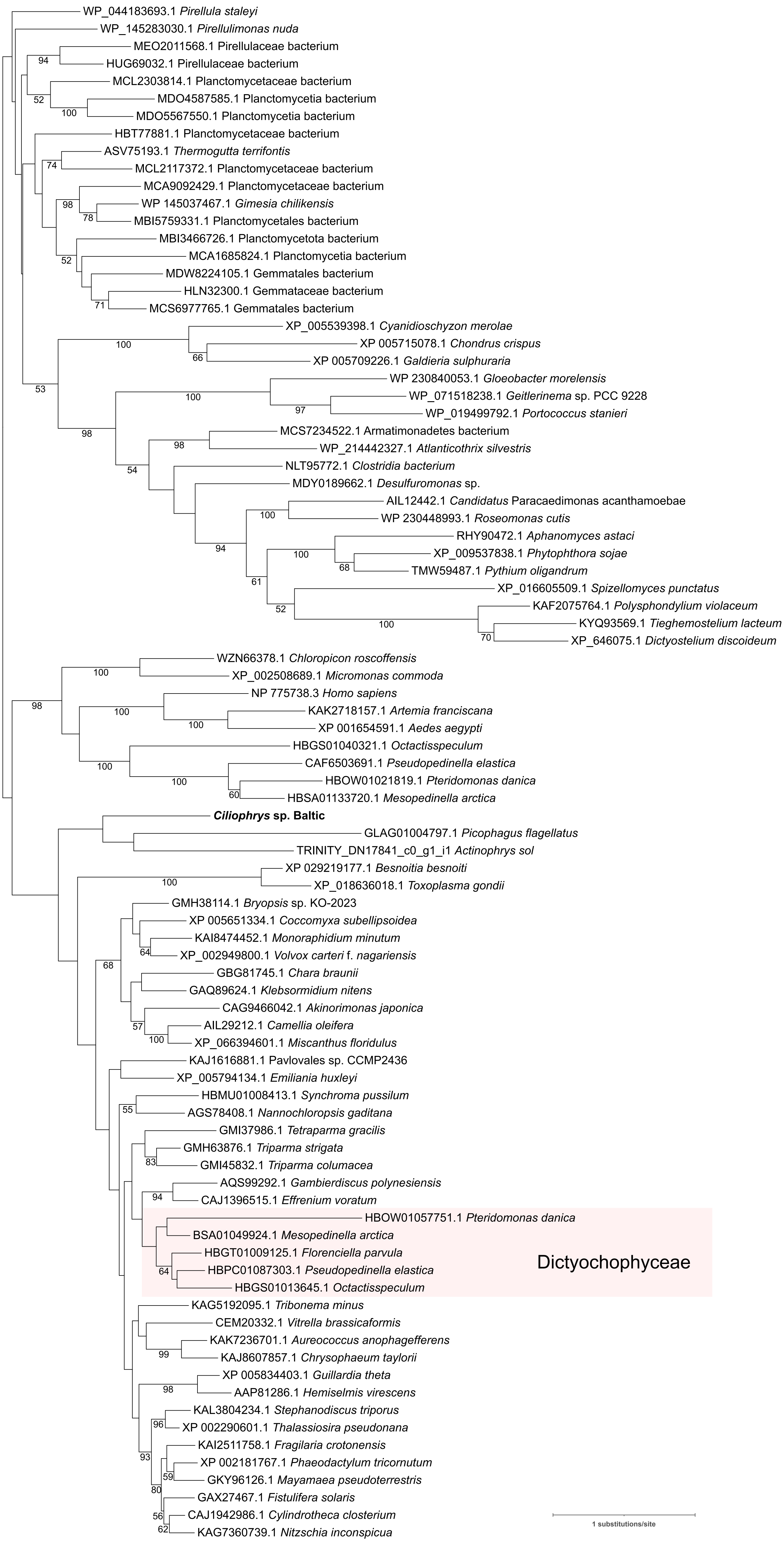


**Fig. S4.** Maximum-likelihood phylogenetic tree of FabD proteins. The final alignment comprised 88 sequences and 406 aligned positions. The phylogenetic tree was inferred using IQ-TREE under the Q.PFAM+R6 substitution model. Numbers at branches indicate bootstrap support values (>50%) calculated from 100 non-parametric bootstrap replicates. The sequence from *Ciliophrys* sp. Baltic is shown in bold (the sequence ID is provided in Table S1). Plastid homologs from other Dictyochophyceae are highlighted with a light pink box.

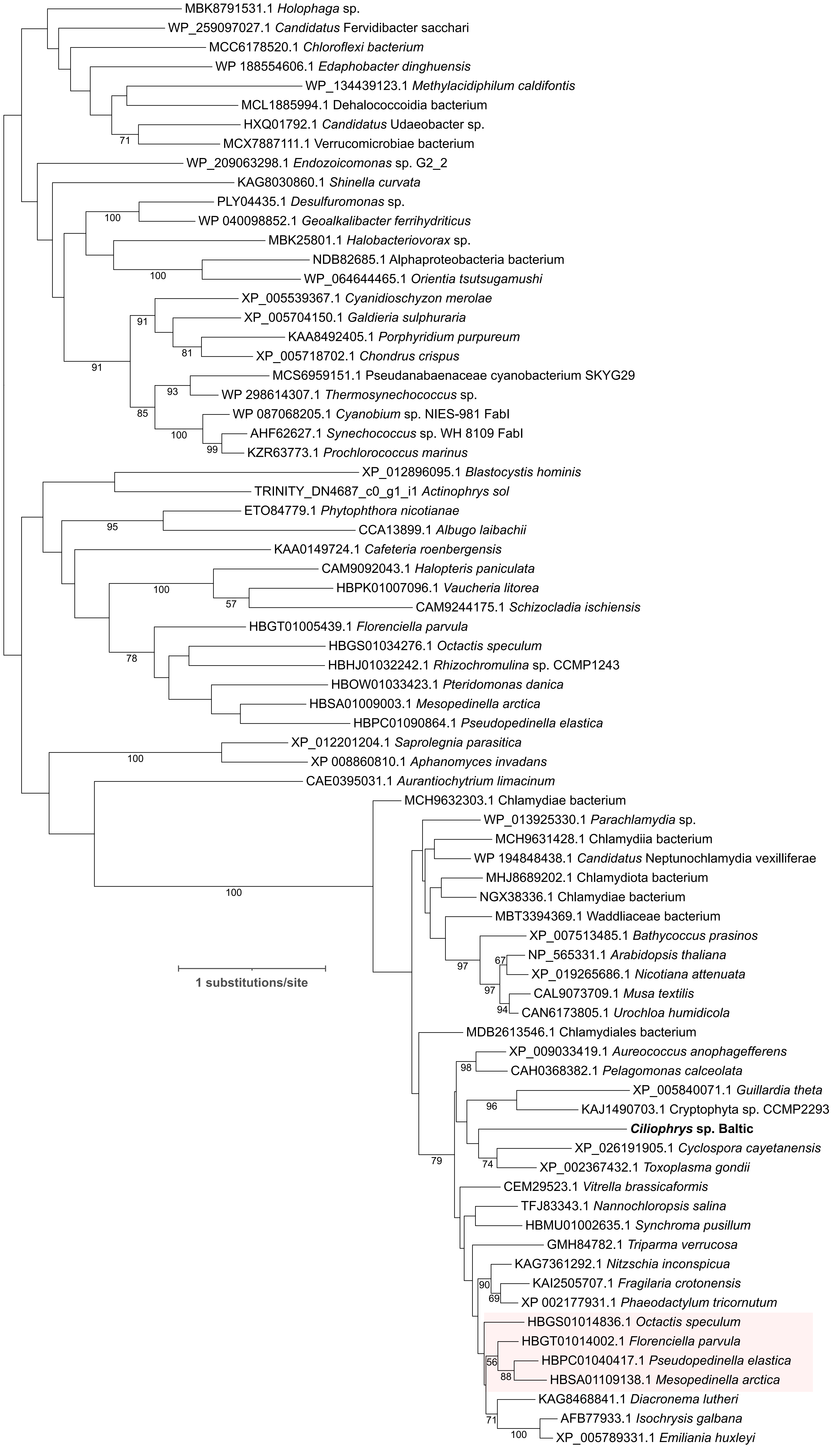


**Fig. S5.** Maximum-likelihood phylogenetic tree of FabI proteins. The final alignment comprised 75 sequences and 371 aligned positions. The phylogenetic tree was inferred using IQ-TREE under the Q.PFAM+R6 substitution model. Numbers at branches indicate bootstrap support values (>50%) calculated from 100 non-parametric bootstrap replicates. The sequence from *Ciliophrys* sp. Baltic is shown in bold (the sequence ID is provided in Table S1). Plastid homologs from other Dictyochophyceae are highlighted with a light pink box.

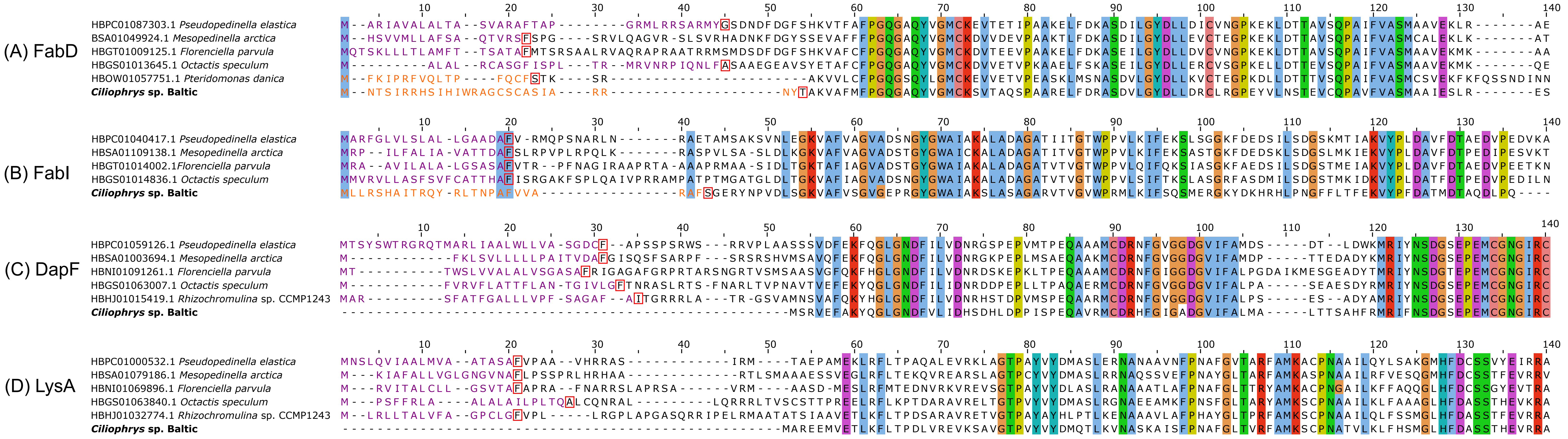


**Fig. S6.** N-terminal sequences of selected plastid-targeted enzymes from dictyochophytes, including their orthologs from *Ciliophrys* sp. Baltic. (A) FabD; (B) FabI; (C) DapF; (D) LysA. Amino acid residues corresponding to N-terminal presequences, namely the signal peptide (in plastid-targeted proteins) and the mitochondrial transfer peptide (in mitochondrion-targeted proteins) as predicted by TargetP-2.0 are highlighted in violet and orange, respectively. The residues immediately following the presequences (i.e., the N-termini exposed upon their cleavage) are boxed in red. Note the absence of any N-terminal presequences in the DapF and LysA proteins from *Ciliophrys* sp. Baltic. Furthermore, as in *Ciliophrys* sp. Baltic, the ortholog of the plastid-targeted FabD in the non-photosynthetic species *Pteridomonas danica* is also predicted to possess a mitochondrial transfer peptide, indicating an independent remodelling of the N-terminal region and a rerouting of the protein to the mitochondrion.

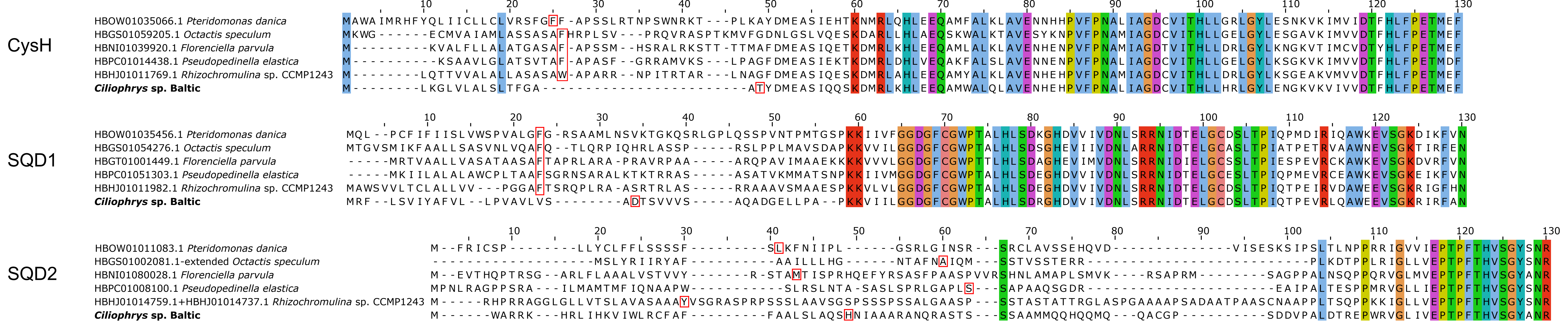

**Fig. S7.** N-terminal sequences of SQDG biosynthesis enzymes. Residues that are immediately downstream of the predicted cleavage site of signal peptide processing peptidase (which removes the N-terminal signal peptide in the ER lumen) are boxed in red.

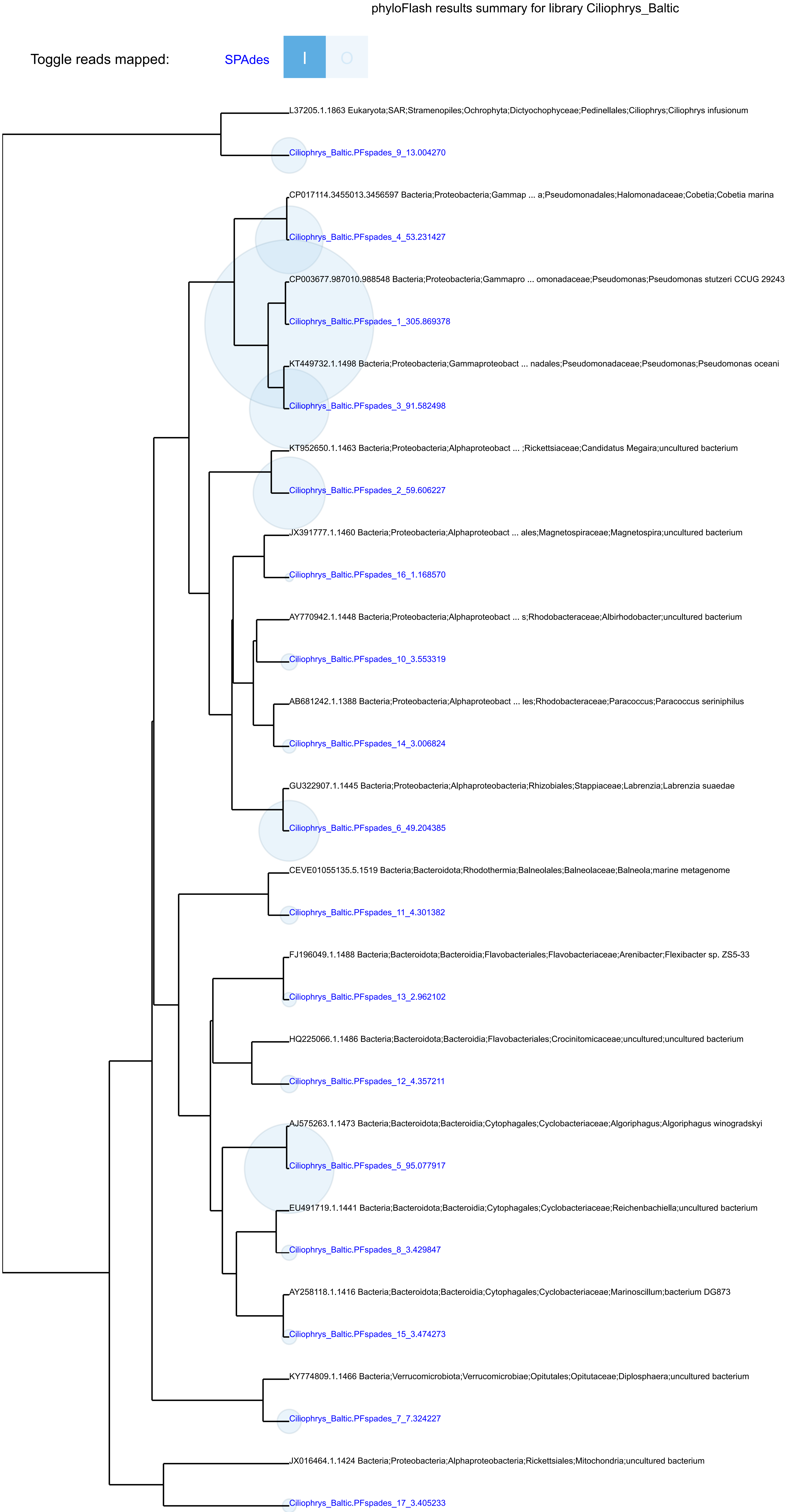


**Fig. S8.** phyloFlash output demonstrating the presence of a variety of bacteria in the (meta)genome data from *Ciliophrys* sp. Baltic.

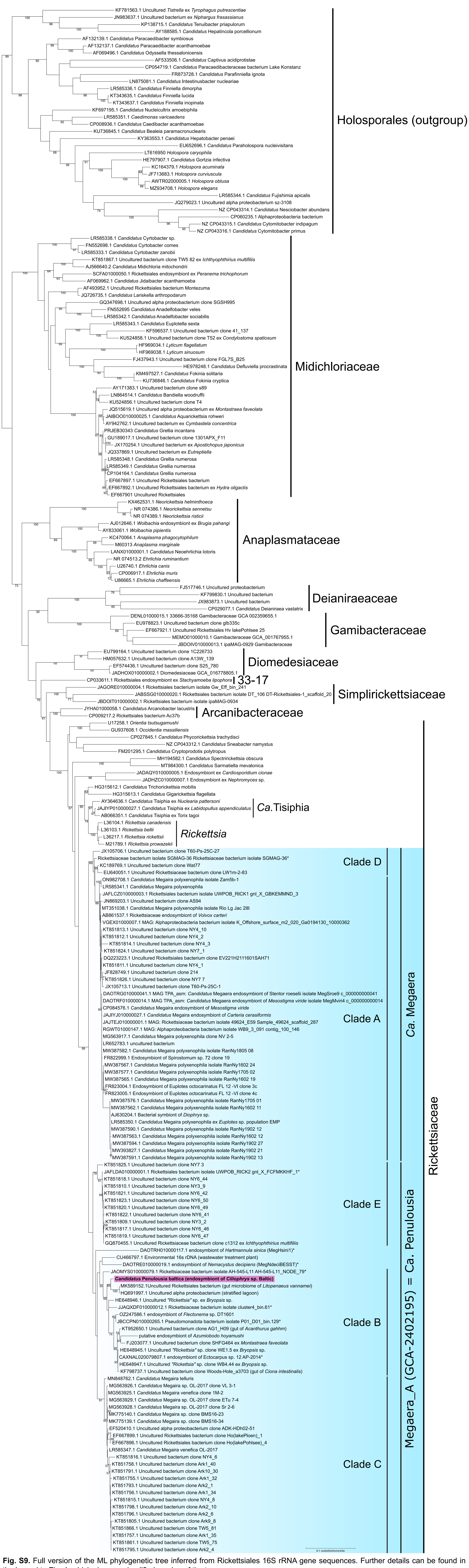

**Fig. S9.** Full version of the ML phylogenetic tree inferred from Rickettsiales 16S rRNA gene sequences. Further details can be found in the legend to Fig. 4, which shows a simplified version of the tree.

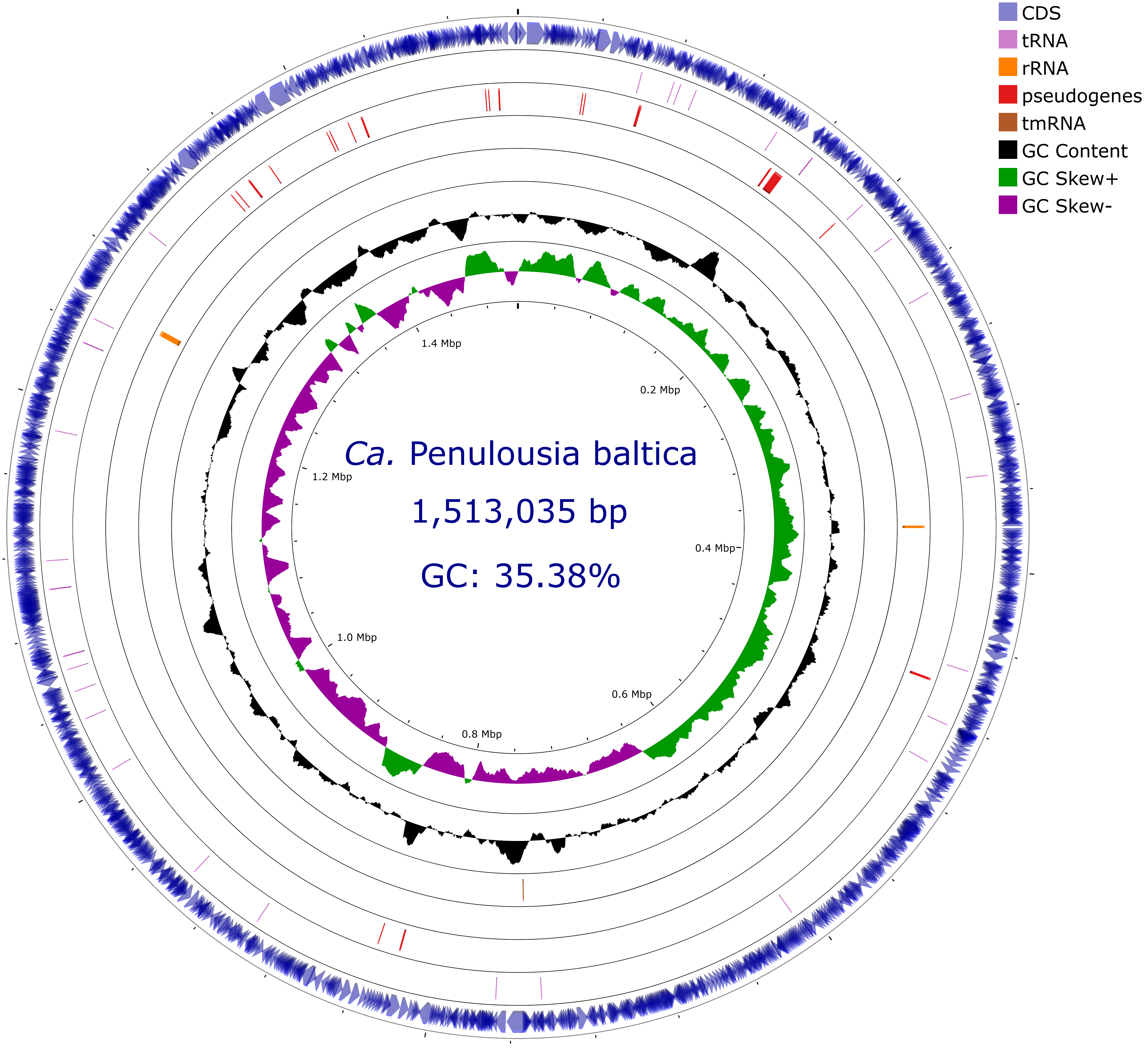


**Fig. S10.** Genome map of *Ca.* Penulousia baltica. The outermost circle shows the positions and orientations of protein-coding genes (CDSs). The second circle indicates tRNA genes, and the third circle shows putative pseudogenes. The fourth and the fifth indicate rRNA and tmRNA genes, respectively. The innermost circles display the GC content (black line) and GC skew (green/purple line), highlighting regions of compositional bias.

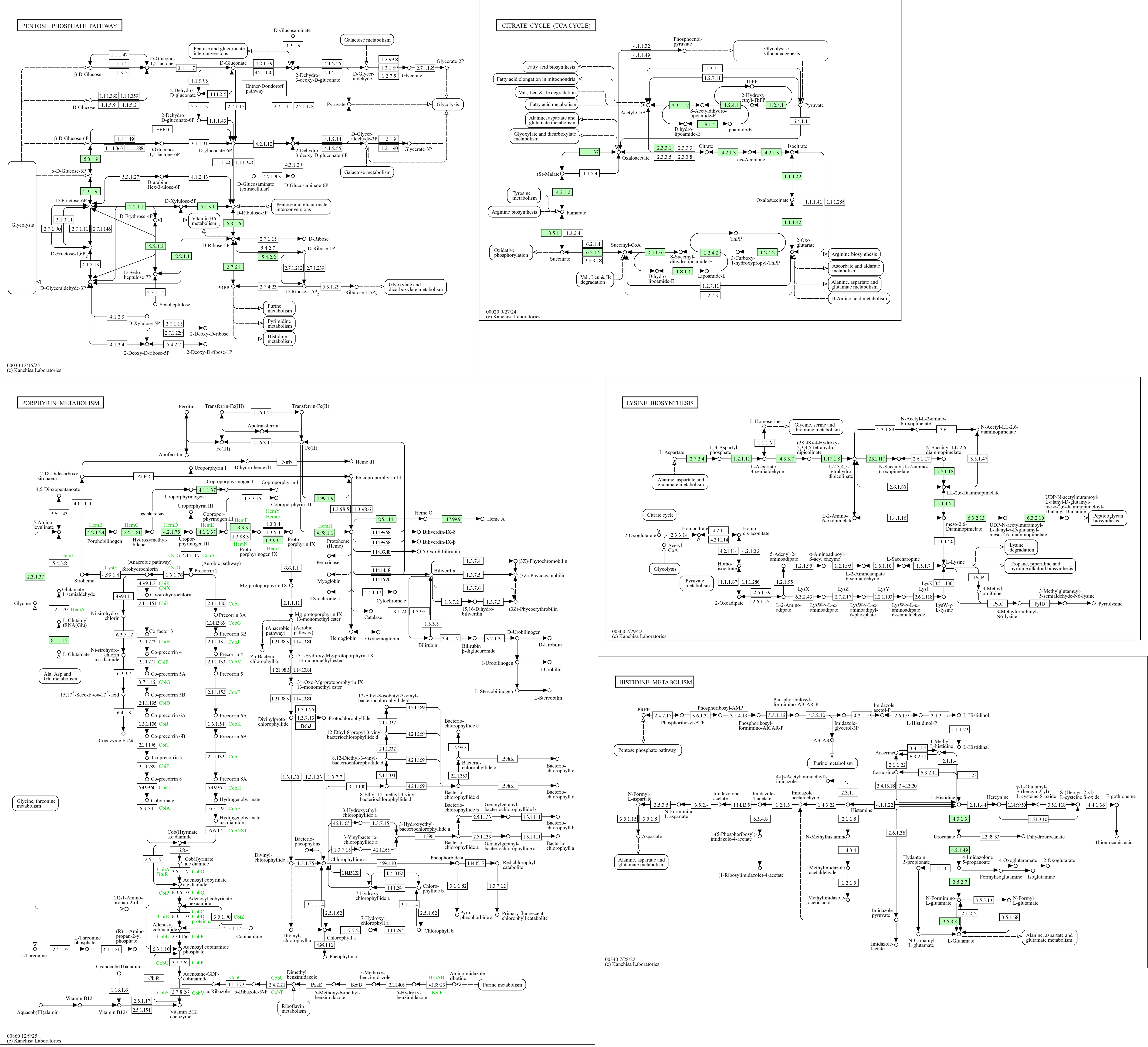


**Fig. S11.** An overview of selected metabolic pathways of *Ca.* Penulousia baltica generated with KEGG Mapper. The schemes indicate the occurrence of particular enzymes in the bacterium (the rectangles in green) as reconstructed using a BlastKOALA search against the KEGG database.

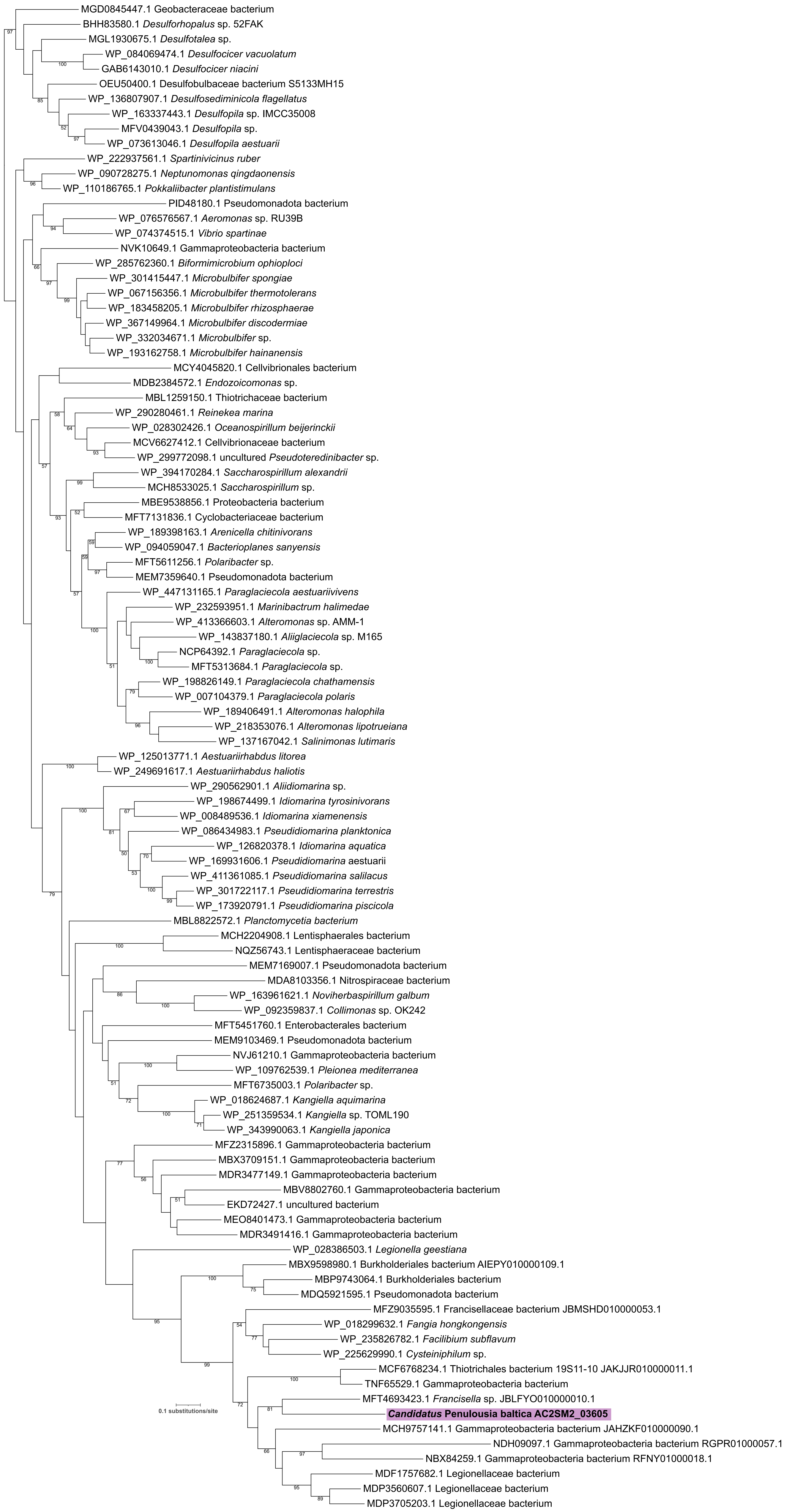


**Fig. S12.** Maximum-likelihood phylogenetic tree of hutH homologs. The final alignment included 101 sequences and 514 positions. The substitution model employed was LG+I+G4 as automatically selected by IQ-TREE. Numbers at branches represent bootstrap values (shown when >50%) calculated from 100 non-parametric bootstrap replicates.

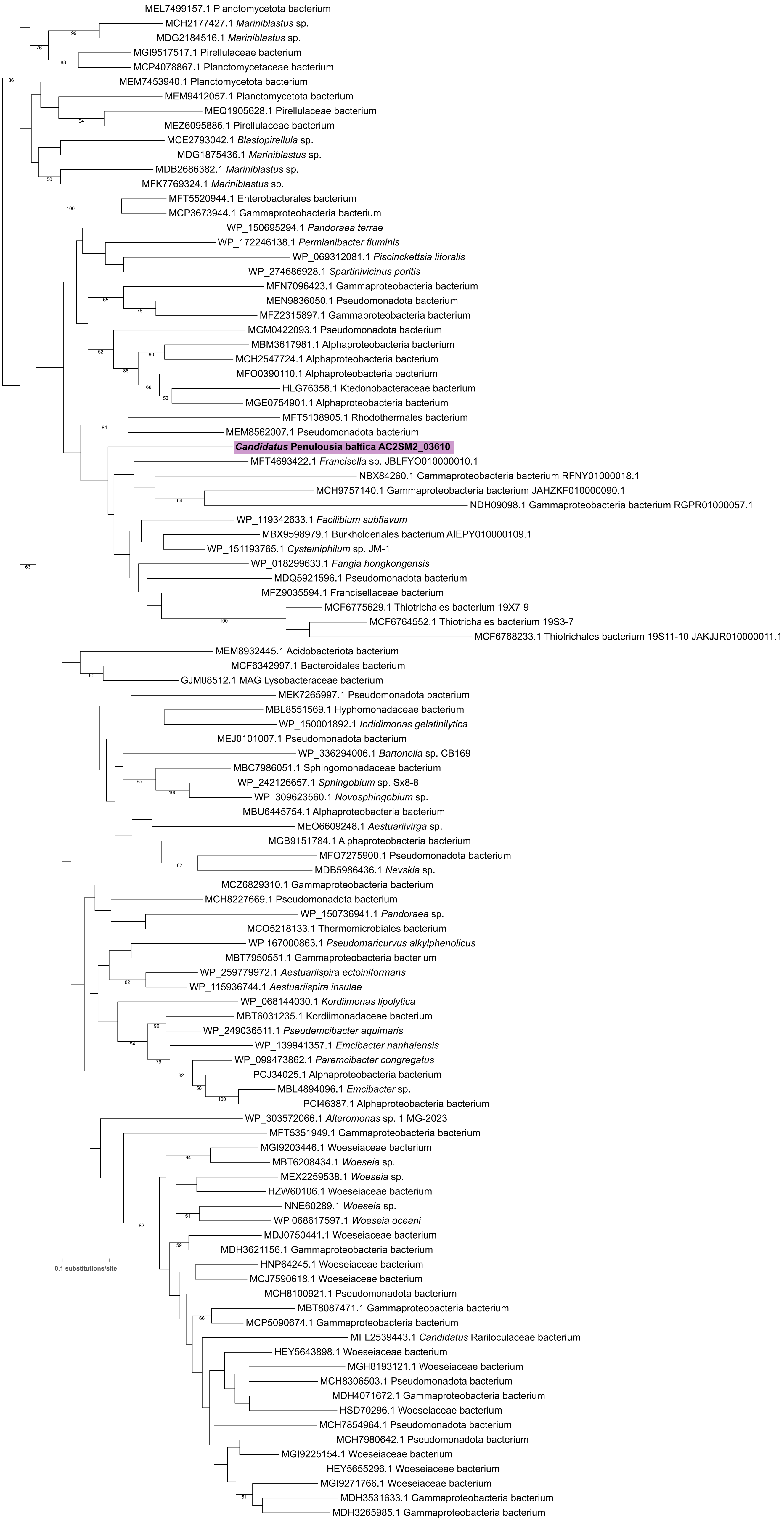


**Fig. S13.** Maximum-likelihood phylogenetic tree of hutU homologs. The final alignment included 104 sequences and 565 positions. The substitution model employed was LG+I+G4 as automatically selected by IQ-TREE. Numbers at branches represent bootstrap values (shown when >50%) calculated from 100 non-parametric bootstrap replicates.

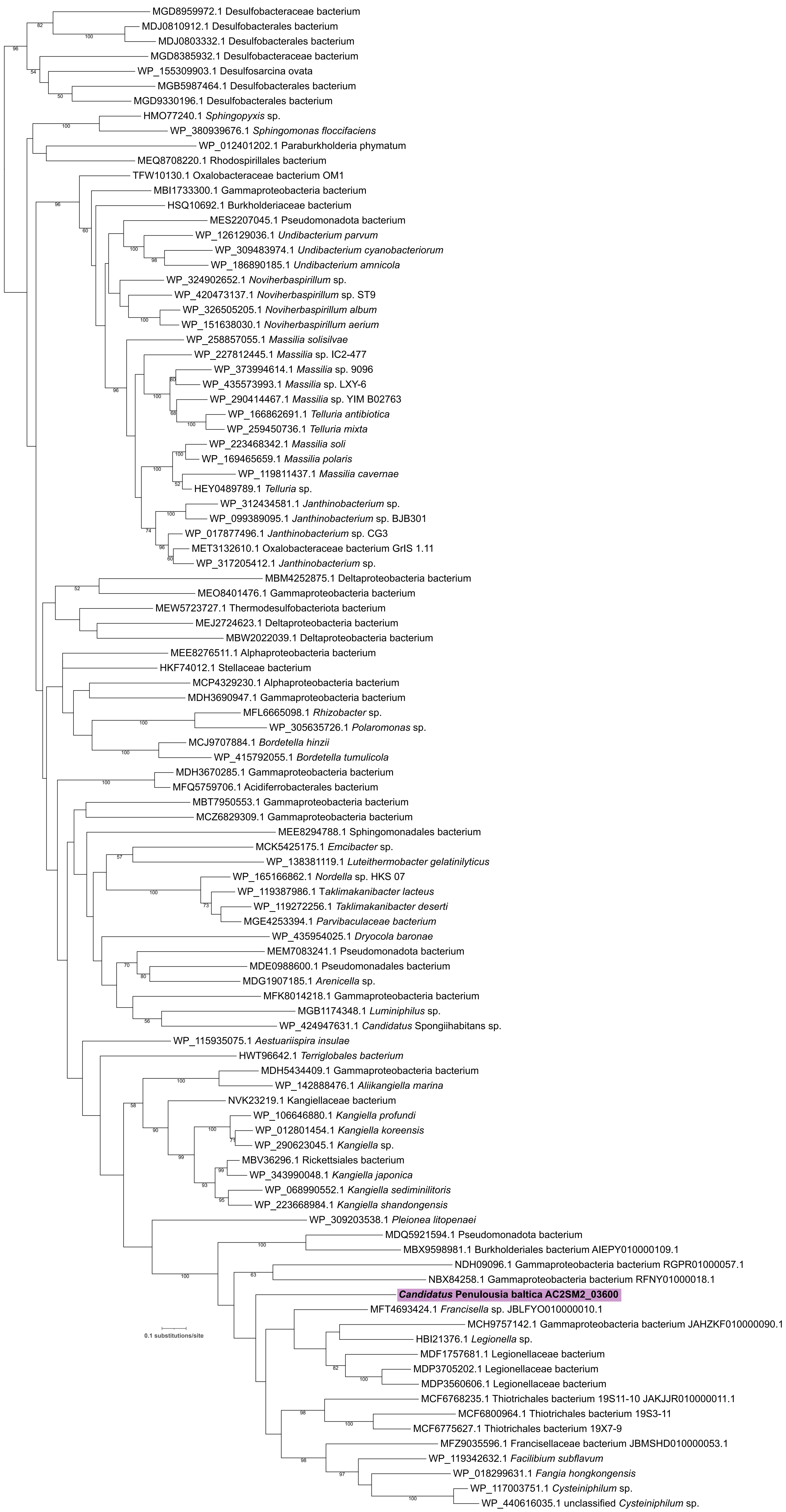


**Fig. S14.** Maximum-likelihood phylogenetic tree of hutI homologs. The final alignment included 101 sequences and 425 positions. The substitution model employed was Q.pfam+I+G4 as automatically selected by IQ-TREE. Numbers at branches represent bootstrap values (shown when >50%) calculated from 100 non-parametric bootstrap replicates.

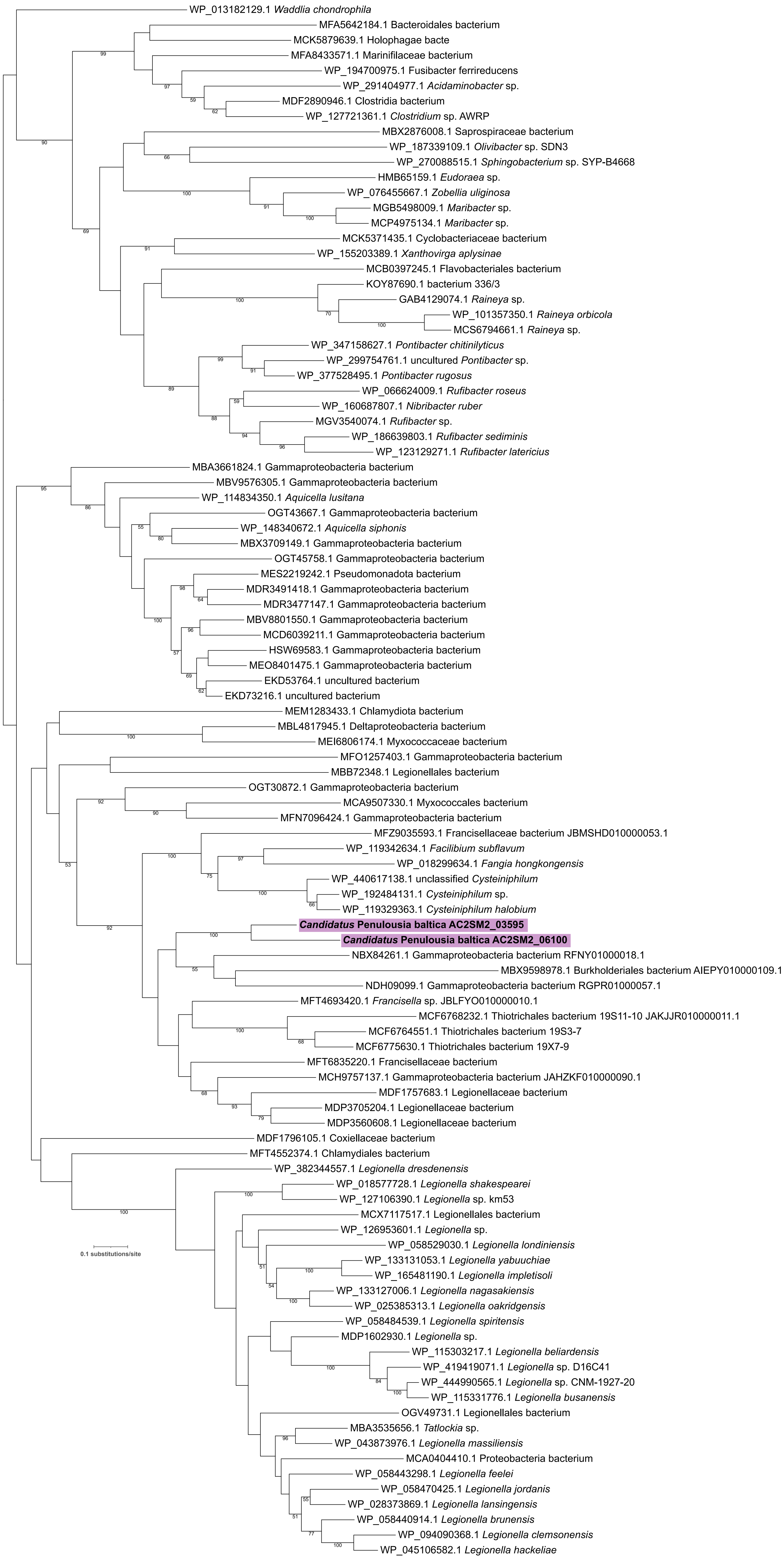


**Fig. S15.** Maximum-likelihood phylogenetic tree of hutG homologs. The final alignment included 102 sequences and 341 positions. The substitution model employed was LG+F+I+G4 as automatically selected by IQ-TREE. Numbers at branches represent bootstrap values (shown when >50%) calculated from 100 non-parametric bootstrap replicates.

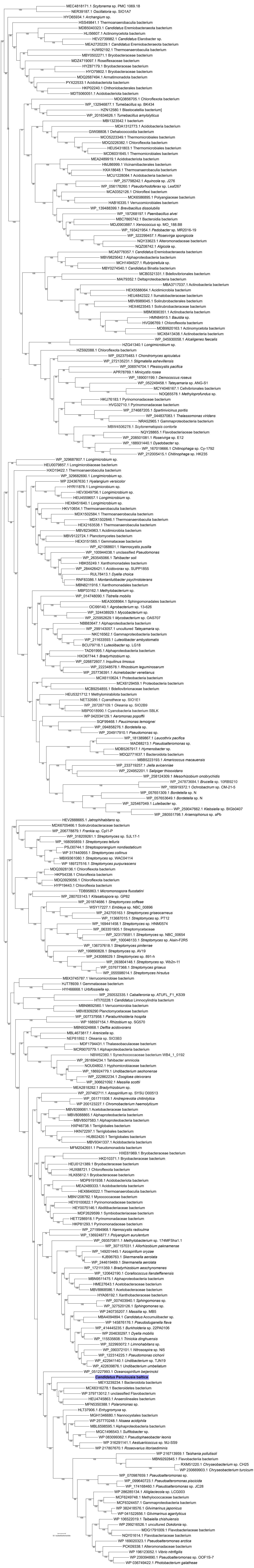

**Fig. S16.** Phylogenetic analysis of LodA/GoxA homologs. The final alignment included 286 sequences and 1042 positions. The substitution model employed was Q.pfam+R10 as automatically selected by IQ-TREE. Numbers at branches represent bootstrap values (shown when >50%) calculated from 10,000 ultrafast bootstrap replicates. The resulting phylogeny did not clearly resolve the donor lineage of the LodA/GoxA gene horizontally transferred into the *Candidatus* Penulousia baltica genome.
